# Rapid chromosomal evolution and polycentromeric drive in sedges and rushes

**DOI:** 10.64898/2026.02.27.708619

**Authors:** James I. McCulloch, Marcela Uliano-Silva, Charlotte J. Wright, Ian R. Henderson, Sam Ebdon, Darwin Tree of Life Consortium, Kamil S. Jaron, Mark Blaxter

## Abstract

The chromosomes of most eukaryotes have a single centromere, a specialised region involved in chromosome partitioning to daughter cells. Inter-chromosomal rearrangement risks generating chromosomes with two centromeres or none, disrupting segregation. Holocentric chromosomes, with centromere function distributed along the chromosome, are hypothesised to better tolerate inter-chromosomal rearrangement. However, evidence linking centromere organisation to rearrangement rate has been lacking. Sedges and rushes (Cyperaceae and Juncaceae) are specifically polycentric: they have several satellite-based centromeres per chromosome, whereas centromere location in “asatellitic” holocentrics is solely defined epigenetically. Using 36 chromosome-level genomes, we quantify extraordinary rearrangement rates and annotate candidate polycentromeres. Under our polycentromeric drive model, we expected satellite sequence, polycentromere architecture, and karyotype to evolve to exploit biased segregation during asymmetric meiosis. We find satellite turnover but also deep sequence conservation. Polycentromeric organisation seems constrained by chromosome size, and rearrangements are more stable when chromosomes have fewer polycentromeres, despite breakpoint regions being enriched for polycentromeres. Notably, we find putative monocentromeres in *Carex myosuroides*, which would be the first evidence for reversion to monocentricity in eukaryotes, and no identifiable polycentromeres in *Cyperus rotundus*, possibly a transition to asatellitic holocentricity. Overall, we demonstrate that this clade is powerful for linking centromere organisation to genome evolution.

## 1. Introduction

Karyotypes are highly variable across eukaryotes, with described haploid chromosome numbers ranging from one^1, 2^ to at least 229^3^. However, the driving forces that generate this variation are still unclear, despite karyotype evolution being implicated in speciation and adaptation^4, 5^. The cyperid clade, predominantly comprising plants in the families Cyperaceae (sedges) and Juncaceae (rushes), is well-suited to investigating these drivers due to the range of haploid numbers represented (*n* = 2–114)^6^. Within this clade, karyotype differs considerably even between species of the same genus, with *Carex* species having *n* = 5–66^7^.

It has been hypothesised that holocentricity – the occurrence of centromeric activity across nearly the entire chromosome length, found in about 20% of eukaryote species^8^ – facilitates chromosome number evolution through inter-chromosomal fusions or fissions. Fission of holocentric chromosomes should not be necessarily deleterious because neither fission product will be acentric, unlike fission products of monocentric chromosomes^9^. Similarly, fusion of monocentric chromosomes yields a dicentric chromosome. In a dicentric chromosome, the two kinetochores may be oriented towards opposite poles, meaning the chromosome is torn apart during anaphase^10^. Holocentric organisms possess mechanisms to ensure uniform orientation. in some cyperids centromere-specific histone H3 (CENH3) localises as a longitudinal groove on condensed chromatids, ensuring mono-orientation^11, 12^. They also often display inverted meiosis, whereby sister chromatids separate first, followed by homologous chromosomes, helping to prevent missegregation of multivalents^13^.

Contrary to these expectations, comparative analysis of insect karyotypes recovered similar rates of chromosome number evolution in monocentric and holocentric taxa^14^. However, this analysis was limited by the number of independent transitions in centromere organisation. Holocentric insects typically have “asatellitic” diffuse centromeric function where kinetochores assemble wherever the chromatin state is permissive^15^. However, in most cyperids studied, centromere-specific satellite arrays interact with CENH3 to recruit kinetochores, similar to the process in monocentrics^12, 16^. We refer to the individual satellite arrays that have centromeric activity as individual polycentromeres, and the chromosomes with many centromere-specific satellite arrays as polycentric. The variation in polycentromere organisation in cyperids allows us to test the relationship between centromere organisation and rearrangement rate. For example, does increased polycentromere density increase the rate of fission, as fission products are less likely to be acentric? Polycentromeres may also directly facilitate rearrangements, acting as common breakpoints, as proposed in the sedge genus *Rhynchospora*^17^.

Polycentromeres and chromosomal rearrangements may be linked by an evolutionary response to centromeric drive. In monocentric species, changes in the length or sequence of centromere-related repeat arrays may bias the inheritance of the chromosome into the gamete when meiosis is asymmetric ^18^. An analogous process – holokinetic drive – has been invoked for asatellitic holocentric species, in which simply changing chromosome length through repeat purging or proliferation, or inter-chromosomal rearrangements (hereafter ‘rearrangements’), could bias inheritance of homologous chromosomes into the gamete^19^. For polycentric species, we synthesise a model of “polycentromeric drive”. Under this model we expect drive to act on satellite sequence, as in classical centromere drive, through impacts on kinetochore stability. We would also expect drive to act on chromosome length, as in holokinetic drive, as this would affect the span of the kinetochore and the number of microtubule attachments. However, in extension, we would also expect drive to act on individual satellite arrays, selecting on total length and density, which may modulate kinetochore robustness or stability. Drive may be especially strong in sedges because meiosis in both sexes is asymmetric^20^.

In this paper, we describe the dynamics of chromosome evolution in the cyperid clade and investigate the relationship between rearrangements and the organisation and position of polycentromeres. Using recently generated cyperid chromosomally-complete genomes, most produced by the Darwin Tree of Life project^21^, we first reconstruct ancestral linkage groups. This allows us to compare rearrangement rates between subclades, analyse the driving forces behind the diversity of karyotypes, and identify fusion and fission breakpoint regions. Secondly, we identify candidate polycentromeric satellite repeats in each species. Using sequence similarity, we classify these repeats and describe their turnover. Finally, by quantifying their organisation and enrichment at rearrangement breakpoints, we explore the relationship between candidate polycentromere arrays and chromosomal evolution.

## 2. Methods

### 2.1 Ancestral linkage group inference

Having selected 38 genome assemblies (Table S1) and reconstructed the phylogeny (Figure S1, Figure S2, detail in Supplementary Text 1), ALGs were inferred using *syngraph* v 0.2.0a^42^ (https://github.com/Obscuromics/syngraph). The inference was run using options -r 2 -s Ananas_comosus -m [1,100]. Our justification for the final value of -m chosen, 33 (Figure S3), is provided in Supplementary Text 2.

### 2.2 Calculation of rearrangement rates

The ALGs inferred at each internal node were used to calculate the rearrangement rate along each internal branch. Given that *syngraph* does not reconstruct gene order, the number of fissions and fusions is likely to be underestimated, but in a consistent way throughout the tree^42^.

We used median divergence time estimates (from TimeTree 5^43^) (Table S2) to date key nodes in our phylogeny for estimation of lineage-average rearrangement (the sum of fusion and/or fission events) rates. The age estimate for the base of the Cyperaceae is the median divergence time of *Schoenus* and *Rhynchospora*^44–47^. The median divergence time of the Juncacae is from the LCA of *Luzula* and *Juncus*^45–56^. The estimate for the base of the *Carex* clade is the median divergence time of *C. myosuroides* and *C. caryophyllea*^50, 57^.

To formally compare rearrangement rates between Cyperaceae and Juncaceae, a Bayesian phylogenetic regression model was used with a negative binomial error distribution, implemented in *brms* v 2.23.0^58–60^. The total number of rearrangements along each lineage, as well as fusions and fissions separately, were modelled as a function of family, incorporating the age of the lineage (time since the basal node of the respective family in our phylogeny) as a log-offset. The rearrangement rates of *Carex* and non-*Carex* Cyperaceae were modelled in the same way.

To investigate whether rearrangement rate may be driven by mating system, heterozygosity, calculated by *k*-mer spectrum analysis using *GenomeScope* 2.0^61^, was used as a proxy for inbreeding. Phylogenetic regression models of rearrangement rate, fission rate, and fusion rate for each species, calculated only along the terminal branch leading to that species, were fitted, with heterozygosity as a predictor, implemented in *brms*^58–60^.

### 2.3 Modelling fission and fusion probability

Four models of fission and fusion probability were tested to investigate the drivers of the observed distribution of tip karyotypes in *Carex*. Each model was based on the number of rearrangements and probability of each rearrangement type (fusion or fission) along internal branches. Random walks, with the same number of steps per branch as rearrangements inferred by syngraph, were simulated with 10,000 trials to produce distributions of the variance of tip karyotypes under the first two models, which either have fusion probability fixed for each branch (“fixed fusion probability” model) or drawn from a Beta distribution fitted using the observed fusion probabilities (“branch-heterogeneous” model). The “fixed fusion probability” model was also run with increased numbers of rearrangements to investigate the impact of underestimating rearrangement rate. We used maximum likelihood estimation to determine whether karyotypes may be shaped by a stabilising force (“stabilising bias” model) or by autocorrelation (“finite-memory heterogeneity” model) acting on fusion probability. These models are explained fully in Supplementary Text 3.

### 2.4 Centromere annotation

*RepeatModeler* v. 2.0.5^62^ was run on all 38 genomes and custom scripts were used to concatenate and cluster consensus FASTA with >95% similarity using *cd-hit-est*^63^. The outputs were passed through *RepeatClassifier* (https://github.com/Dfam-consortium/RepeatModeler/blob/master/RepeatClassifier). The library was then used for *RepeatMasker*^64^. Genes were annotated using *Helixer* v 0.3.5^65, 66^. Centromere annotation was performed using *CAP* (https://github.com/vlothec/CAP/tree/main) using the locations of genes and repeats identified by *Helixer* and *RepeatMasker* in gff3 format. Visual inspection of satellite distribution using *CAP* output plots, which can be found at https://github.com/Obscuromics/cyperids/upload/main/data/CAP, with some input from its centromere probability metric, was used to identify candidate centromere-associated repeats (see Supplementary Text 4). The consensus sequence for each candidate satellite for each species was retrieved from the *CAP* output. To classify the satellites into families, consensus sequences from each species were decomposed into 8-mers and Jaccard similarity was calculated. Satellites were classified into the same family if Jaccard similarity was > 0.1 to any other member of the family. This threshold is sufficient to group the two previously-classified *Tyba* satellites^12^ in our dataset, which have a Jaccard similarity of 0.138. Setting this threshold lower risks including unrelated repeats in the same family. We chose *k* = 8 to account for the shorter satellite monomers in our dataset. The complete similarity matrix is available at https://github.com/Obscuromics/cyperids/tree/main/data.

We used *BLASTn-short* (*BLAST* v 2.15.0) to investigate whether satellite families were present in species other than those in which they were classified as a candidate polycentromeric satellite^67^. We compared the candidate polycentromeric satellite consensus sequences from each species against a database of all satellites identified by *TRASH*^68^, using an E-value threshold of 1e-3 and considering satellites to have shared ancestry if the alignment covered ≥ 40% of the shorter sequence. The repeat *Mateo* in *Carex myosuroides* has a 34-bp stretch of AT dinucleotide repeats which resulted in hits to likely unrelated satellites; in *Carex littledalei*, this region has only 10 bp of AT dinucleotide repeats, so *Mateo* hits were disregarded if they were only to *Mateo* in *C. myosuroides* and not also *C. littledalei*.

### 2.5 Characterising the effect of candidate polycentromeres on the TE and coding sequence landscape

To relate the position of candidate polycentromeres to the landscape of transposable elements and genes we divided chromosomal scaffolds into 5 kb windows. The presence or absence of a candidate polycentromere-associated satellite within each window was recorded. We chose this window approach to minimise the risk of TE insertions erroneously dividing candidate centromeric arrays. Some particularly large TE insertions may still pose a problem, but increasing the window size further would unacceptably inflate array size given that the smallest candidate centromeric arrays average approximately 5 kb. For each non-polycentromeric window, we calculated the distance to the nearest candidate polycentromeric array, as well as whether the window was overlapped by a gene’s coding sequence (CDS, predicted by *Helixer*^65, 66^) and/or by a transposable element (identified by *RepeatModeler*^62^). We then fitted a binomial generalised linear model of CDS overlap and TE overlap separately, predicted by log distance to the nearest centromeric array. We removed data from the windows immediately next to the centromere and enforced a maximum distance of half the mean inter-array gap, calculated as described below.

### 2.6 Relating candidate polycentromeric distribution to rearrangement rate and chromosome size

To relate the distribution of candidate polycentromeres to rearrangement rate and chromosome size, we calculated the mean and median candidate polycentromeric array length and mean and median inter-array gap length using our measurements on 5 kb windows. We also calculated polycentromeric share (the summed length of arrays divided by the length of the non-polycentromeric genome) and array density (the number of arrays divided by the length of the non-polycentromeric genome). We used windows rather than exact array lengths because polycentromere-related arrays could be punctuated by non-satellite DNA. We defined share and density relative to the length of the non-centromeric sequence to avoid the spurious inflation of correlations given that the length and density of arrays would otherwise contribute to overall chromosome length.

For chromosome fusion, fission, and overall rearrangement rate, nested Bayesian phylogenetic regression models were constructed, implemented in *brms* v 2.23.0^58–60^. Rates were modelled using a lognormal error distribution. Given their close relatedness, *Bolboschoenus planiculmis* and *x Bolboschoenoplectus mariqueter* were collapsed into one observation, incorporating the mean of the polycentromere organisation variables, and the rearrangement rate along the branch terminating in their last common ancestor (LCA).

Taking overall chromosomal fusion and fission rate and genome-wide averages of polycentromere distribution variables may mask patterns within a genome. We first fitted phylogenetic regression models of chromosome length, which was log-transformed and modelled with a Gaussian error distribution. Then, to investigate associations between candidate polycentromere organisation and rearrangements on a within-species, chromosomal level, we used the *syngraph* inference to record whether each chromosome was a product of fusion, fission, or both since the species’ parent node. We then modelled each of the four variables of candidate polycentromere organisation with phylogenetic linear regressions and a Gaussian error distribution, using fusion history, fission history, and log-transformed chromosome size as predictors.

### 2.7 Testing for candidate polycentromere enrichment at breakpoints

To test whether polycentromeres may directly facilitate fission and fusion events, we identified breakpoint regions in 8 pairs of sister taxa and compared the candidate polycentromere share within and outside of these regions using Fisher’s exact tests. To identify the regions, we adapted the method of Zhang *et al.*^17^, using *minimap2* v. 2.28 for reciprocal whole-genome alignments of these species pairs (-cx asm10)^69^. Where this resulted in an alignment of one “target” chromosome against two or more “query” chromosomes, the region on the target chromosome where the aligned query chromosome switches was defined as a candidate breakpoint region. Such regions could represent fusions in the genome of the “target” chromosome, fissions in the genome of the “query” chromosome, or alignment errors. To confirm and classify the breakpoint regions, we used the synteny of BUSCOs between the two genomes, and their assignment to ALGs in the last common ancestor.

## 3. Results

### 3.1 Rates of inter-chromosomal rearrangement in the cyperid clade

To investigate the dynamics of chromosome rearrangement across the cyperids, we used chromosome-level genome assemblies from 36 diploid species, 31 from the sedge family (Cyperaceae; all Cyperaceae are considered to be polycentric^70^), and five from the rush family (Juncaceae, including two polycentric *Luzula* and three monocentric *Juncus* species). The assemblies were of high completeness (a mean of 90.7% BUSCOs complete and single copy (s.d. = 2.82%)). Our phylogeny constructed with these orthologues concurs with that of Larridon *et al.*^71^ except that *Trichophorum* was resolved as sister to *Eleocharis* and *Scirpus*, rather than to *Carex* (Figure 1A).

**Figure 1.**
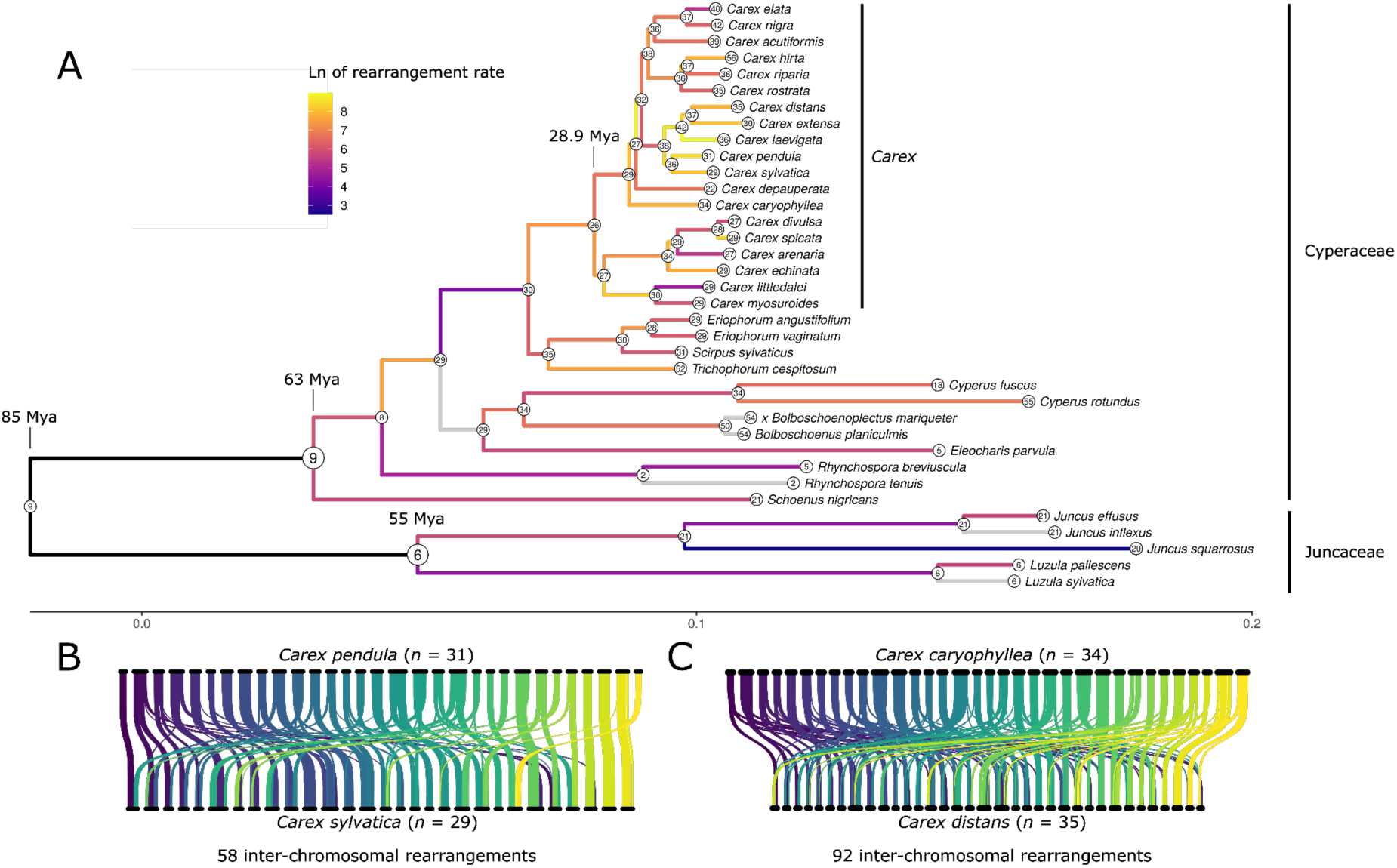
Dynamic karyotypic changes in cyperid species. **A**. Phylogeny of cyperid species analysed. The number of ALGs at the internal nodes and the karyotypes at the tip nodes are represented by the numbered circles. Grey branches lack rearrangements. Rearrangements were not inferred for the black branches given the unreliability of the ALG inference at the base of the cyperid clade. **B**. A ribbon plot showing the macrosynteny of *Carex pendula* and *C. sylvatica*, two sister taxa. Ribbons connect groups of at least 10 syntenic markers, illustrating how roughly similar chromosome numbers in closely related species hide a plethora of independent fissions and fusions. **C**. The macrosynteny of *C. caryophyllea* and *C. distans*, which are less closely related than *C. pendula* and *C. sylvatica* and, accordingly, are even more rearranged despite similar karyotypes.

Overall, chromosomal fissions were more common than fusions, and the rearrangement rate was higher in Cyperaceae than Juncaceae, and highest in *Carex* (Figure 1A, Figure S4, Figure S5). For the Cyperaceae, the median rate across all tips since the basal node was 1.175 inter-chromosomal rearrangements per million years (r/My) (s.d. = 0.445, n = 32 non-independent lineages), with 0.444 fusions per million years (fu/My) (s.d. = 0.202) and 0.762 fissions per million years (fi/My) (s.d. = 0.274). For the Juncaceae, the median rate estimate was 0.291 r/My (s.d. = 0.106, n = 5 non-independent lineages), 0.073 fu/My (s.d. = 0.035), and 0.273 fi/My (s.d. = 0.123).

Branches with particularly high rates of inter-chromosomal rearrangement were found within *Carex* in our phylogeny. The median temporal rate of inter-chromosomal rearrangement for species of *Carex* was estimated as 1.31 r/My (s.d. = 0.718, n = 19 non-independent lineages), with 0.588 fu/My (s.d. = 0.338) and 0.623 fi/My (s.d. = 0.364). The branch with the highest rate of rearrangements was between *Carex laevigata* and its LCA with *C. distans* and *C. extensa*, along which 60 rearrangements are inferred to have occurred (26 fissions and 34 fusions). The slope estimate of the phylogenetic regression model of rearrangement rate as a function of family was 0.497, suggesting that the rearrangement rate of the Juncaceae in our dataset is about half that of the Cyperaceae, however the 95% credible interval (95% CI) overlapped 1 (0.172 - 1.57), and was therefore not significant, likely due to the small sample size for the Juncaceae (n = 5). A similar pattern was found when fusions and fissions were modelled separately, with the fusion rate of Juncaceae estimated to be 0.343 that of Cyperaceae (95% CI: 0.110 - 1.23) and the fission rate estimated to be 0.631 that of Cyperaceae (95% CI: 0.190 - 2.25). The rearrangement rate for non-*Carex* genera was estimated to be 0.60 that of *Carex* (95% CI 0.346 - 1.06). The fusion rate of non*-Carex* genera was estimated to be 0.512 that of *Carex*, with a 95% CI of (0.320 - 0.860). The fission rate of non-*Carex* genera was estimated to be 0.650, but with a 95% CI of (0.330 - 1.32).

It is also pertinent to compare the rearrangement rates of the polycentric genus *Luzula* and monocentric *Juncus*. *Juncus effusus*, *J. inflexus*, and *J. squarrosus* have accumulated 23, 19 and 16 inter-chromosomal rearrangements since the two genera diverged 55 Mya, respectively, while *Luzula pallescens* and *L. sylvatica* have accumulated 12 and 8, the opposite pattern to that expected. This analysis is limited by the small number of monocentric Juncaceae in our dataset.

Inbreeding might favour inter-chromosomal rearrangements by reducing the incidence of structural heterozygosity, which can cause problems with segregation^13^. Additionally, inbreeding can facilitate the fixation of mildly deleterious mutations through drift. Phylogenetic regression models did show a trend of decreasing rearrangement rate with increasing sequence heterozygosity, a proxy for the level of inbreeding, though these relationships were also not statistically significant (overall rearrangement rate: -0.68, 95% CI:-1.52 – 0.19; fusion rate: -0.60, 95% CI: -1.51 – 0.34; fission rate: -0.27, 95% CI: -1.19 – 0.66).

### 3.2 No karyotype-stabilising forces are needed to explain patterns of rearrangement in *Carex*

What forces drive chromosome number evolution? In monocentric taxa, the number of available centromeres strongly patterns chromosome number evolution, but this constraint may be largely relaxed in holocentric taxa. The *Carex* clade has high rearrangement rates and variance (var = 55.813) in tip karyotypes. We developed four models (described in SupplementaryText 2) which differed in the per-branch probability of each rearrangement being a fusion (*pi*) or a fission (1 - *p_i_*).

The “fixed fusion probability” model, in which rearrangements on every branch have an equal probability of being a fusion, predicted a lower variance in chromosome numbers, though not low enough to reject the model (empirical quantile = 0.866, Figure S7). The “branch-heterogeneous” model, in which per-branch fusion probability is drawn from a Beta distribution fitted to observed fusion probabilities (Figure S6), did not result in a distribution with greater density at the observed variance (empirical quantile = 0.128, Figure S7).

As the number of rearrangements may be an underestimate, simulations were also run for the “fixed fusion probability” model where the number of rearrangements on each branch is multiplied by a factor *m*. Maximum likelihood estimation using log of variance in tip chromosome number was employed to estimate a value for the rearrangement rate multiplier *m* resulting in an estimate of 1.611058. Simulations of karyotype evolution run with values of *m* of 1, 1.611058, and 2 showed that rearrangement rate underestimation may affect predictions of tip karyotype variance (Figure S8).

Other factors could be shaping the observed variance in tip chromosome number. The “stabilising bias” model increases fusion probability following a fission-driven increase in chromosome number, and the inverse, constraining karyotype around an optimum value. Such a value could be derived from an optimum recombination rate, balancing mitigating Hill-Robertson interference and maintaining linkage between co-adapted alleles, as there are thought to be typically one or two crossovers per chromosome^72, 73^. However, maximum likelihood estimation of the tuning parameter *ɣ*, which determines the strength of fusion probability modulation in response to chromosome number at the parent node, using optimal chromosome numbers of 26 or 31 (the *Carex* ALG number and median tip chromosome count respectively) gave greatest support to *ɣ* = 0. Thus the “stabilising bias” model does not provide a better explanation for observed fusion and fission proportions than binomial sampling variation when *p* is fixed. The “finite-memory heterogeneity” model assumes that the fusion probability of a branch tends to the fusion probability of its parent branch, with the strength of this tendency modulated by *⍴*. Maximum likelihood estimation provided an estimate of *⍴* = 0, suggesting autocorrelation along lineages also cannot better explain observed fusion and fission proportions than binomial sampling variation.

Overall, these models suggest that variation in chromosome number in *Carex* is largely consistent with stochastic variation around a fixed fusion probability, as variable fusion probabilities, stabilising bias, or autocorrelation do not substantially improve model fit.

### 3.3 Cyperids have diverse candidate polycentromeric satellites

We identified candidate satellite arrays associated with polycentromeres for all diploid species in our dataset (Table S3) except for *Eleocharis parvula*, for which the relative abundance of satellite DNA precluded confident identification of satellites likely to be specifically associated with centromeric activity, using *CAP* (https://github.com/vlothec/CAP), which integrates repeat annotation, gene annotation, GC content, and measures of sequence predictability. . The candidate centromere-associated satellites were classified using a threshold of at least 0.1 Jaccard similarity into 25 families across the 35 species, implying frequent polycentromeric turnover, including between species of the same genus (Figure 2, Figure S9, Table S4, Supplementary Text 4). We recovered all the previously known cyperid polycentromeric repeat families, including the *Tyba* satellite family of *Rhynchospora* centromeres^12^, satellites *Lusy1* and *Lusy2* of *Luzula sylvatica*^74^ (classified in the multispecies families *Isaac* and *Jerry* along with satellites from *Luzula pallescens*), and *JefSAT1*, *JefSAT2*, and *JefSAT3* of *Juncus effusus*^75^ (classified in the families *Amelia*, *Simms*, and *Cheyenne* along with satellites from *Juncus inflexus*). Strikingly the family *Carol* was found in 13 species of *Carex*, with a high average 8-mer Jaccard similarity (0.880). Some putative polycentromeric repeats were conserved across genera. For example candidate centromeric satellites in the *Sandra* family were found in some *Carex*, *Eriophorum*, and *Scirpus*. Across species, there is no overall pattern discernible when comparing the GC content of the centromeric fraction to the genome as a whole (Figure S10).

**Figure 2.**
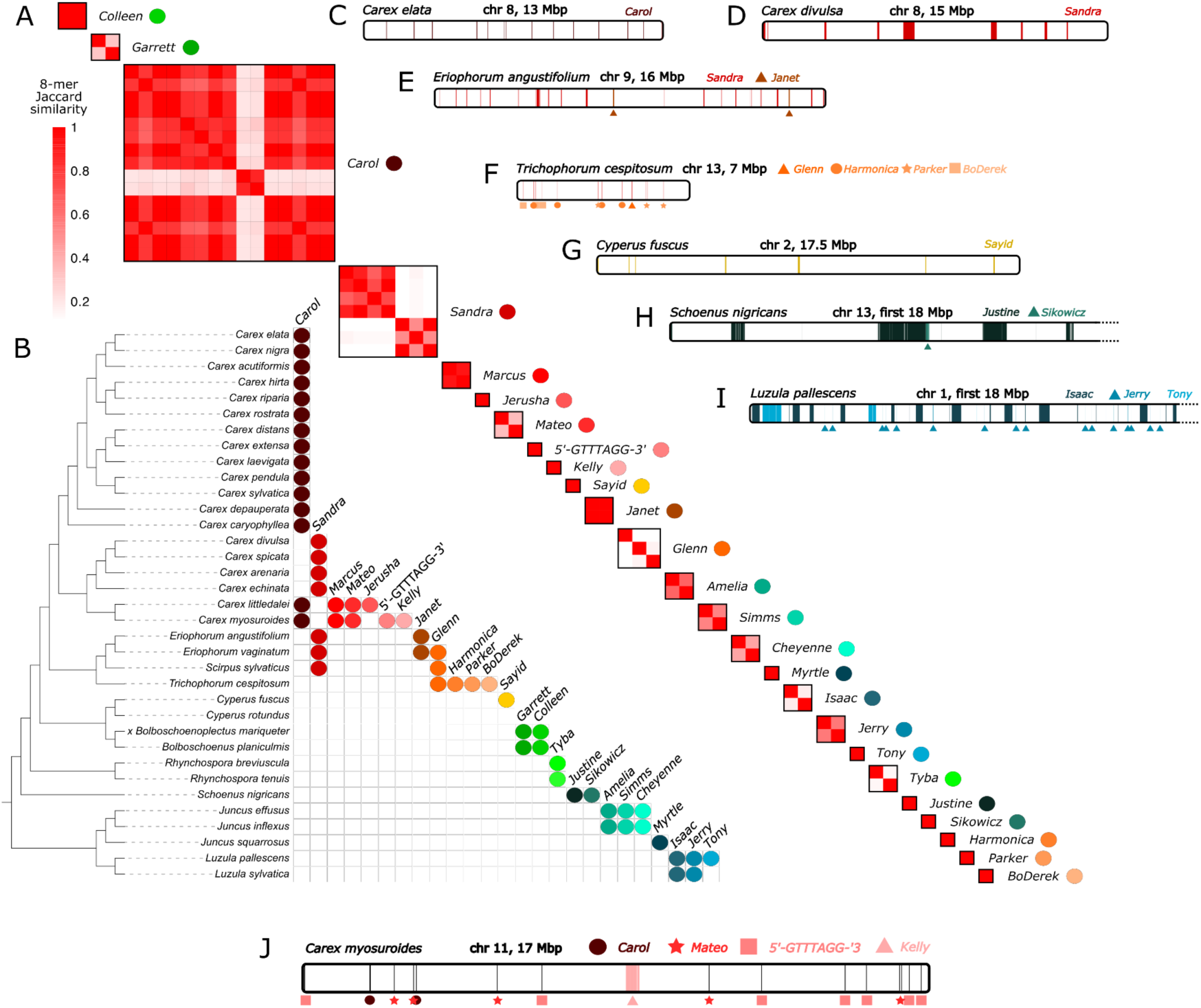
Polycentromeric turnover in the cyperid clade. The candidate polycentromere-associated satellites vary within the cyperid clade. **A**. The heatmap shows families of candidate polycentromeric satellites, defined as repeats with > 0.1 Jaccard similarity using 8-mers. A version with species labels is presented in Figure S10. **B**. The distribution of each family on our cyperid phylogeny. The phylogeny was visualised using iTOL (https://itol.embl.de/)^40^. **C-J**. The arrays of the candidate polycentromere-associated satellites on chromosomes of a selection of species are illustrated beneath to demonstrate the wide variation in the organisation of candidate polycentromeres across the clade. Colours indicate different families of satellite repeats. **J** shows an example chromosome of *Carex myosuroides* which harbours a potential neo-monocentromere underpinned by *Kelly* repeats.

Most chromosomes of *Cyperus rotundus* lacked satellite repeats away from the telomeres. There was no evidence of transposable element-based centromeres. It is possible that this species displays asatellitic holocentricity as seen in some insects^15^.

We identified one potential case of transition from polycentricity to monocentricity. In *Carex myosuroides*, a single array of the 177 bp satellite *Kelly*, ranging in length from 16 kb to 251 kb (median 69 kb), was found near the centre of eight of the 29 chromosomes (Figure 2J). The chromosomes with the longest arrays (chromosomes 2, 9, and 11) showed a noticeable trend of transposable element enrichment and a reduction in gene density towards these possible neo-monocentromeres, akin to the pattern seen around canonical monocentromeres where there is low meiotic crossover^76^. Statistical significance was confirmed by the summary statistics of binomial GLMs of TE density and CDS density with distance to the *Kelly* arrays (TE density: slope = -5.270e-05, p = 0.0198; CDS density: slope = 1.524e-04, p = 1.41e-11).

To explore the origins of candidate centromeric satellites we performed nucleotide similarity searches with the satellite consensus sequences against all satellites for each species, identifying several cases where the satellites were present but not implicated in polycentromere function. (Figure S11, Table S5). The most extreme example was *Parker*, which appeared to be an ancestral satellite for all Cyperaceae in our tree, yet only had polycentromeric signatures in *Trichophorum cespitosum*. Repeat *Kelly* occurred in six species of *Carex*, but only formed monocentromere-like arrays in *C. myosuroides*.

### 3.4 Candidate polycentromere positioning correlates with the TE and coding sequence landscape of the chromosome

We reasoned that we would not see a marked reduction in gene density and enrichment of transposable elements near candidate polycentromeres as is typically seen close to monocentromeres^77^ because having multiple centromeres per chromosome might select against the accumulation of pericentromeric heterochromatin. However, binomial generalised linear models revealed comparable magnitudes of reduced CDS density and TE enrichment with decreasing distance to the nearest centromere in polycentromeric species as in the monocentric *Juncus* species (Figure S12) albeit on shorter scales. Nine polycentric sedge species had steeper slopes of TE enrichment than all three monocentric *Juncus* species, and four polycentric sedge species had steeper slopes of reduction in CDS density, though this comparison is limited by the few monocentric species available. All but seven sedge species had a significant TE enrichment slope, and all but three had a significant reduction in CDS density slope. Only two (*Carex littledalei* and *Cyperus fuscus*) had neither slope significant. *Rhynchospora breviuscula* and *R. tenuis* were outliers; both had a significant increase in TE density and decrease in CDS density away from the nearest candidate polycentromere. Overall, however, the polycentromeres we identified seem to have a similar impact on the sequence landscape as do monocentromeres.

### 3.5 Inter-specific rearrangement rate is partly explained by candidate polycentromere organisation

Given the variation in candidate polycentromere array structure and distribution (Figure 3A), we asked whether this might explain the inter-specific variation in rearrangement rates in the cyperid clade. To investigate this, we related polycentromeric share of the genome, polycentromere density, array length, and inter-array gap length to the fusion and fission rate estimates, only from the tip branches due to polycentromeric turnover (Figure 3B, Table S7).

**Figure 3.**
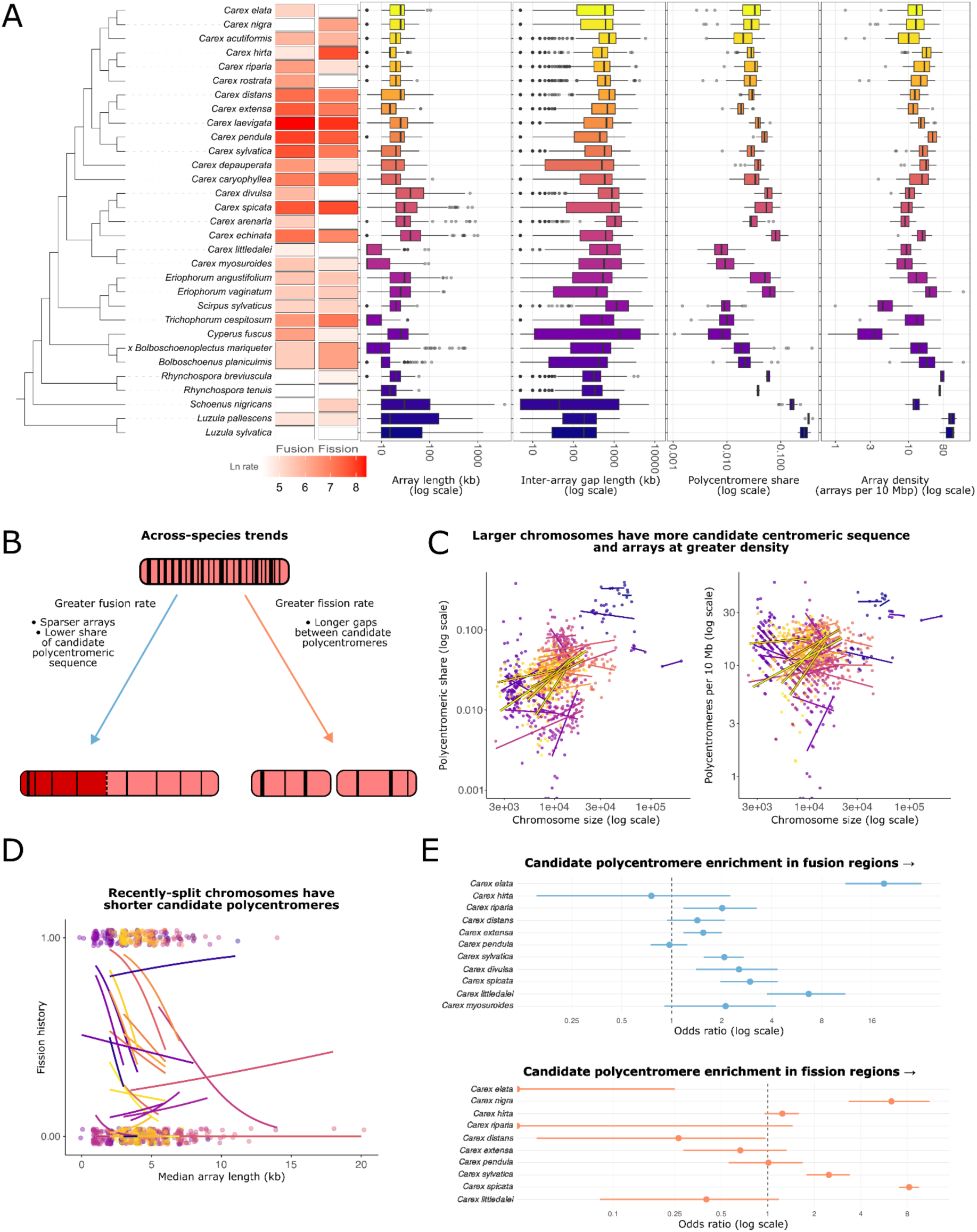
Polycentromeres and inter-chromosomal rearrangements. **A**. A phylogeny of the cyperid species for which candidate polycentromeric arrays could be identified, showing for each species the values of four polycentromeric arrangement variables: array length, inter-array gap length, polycentromeric share of the chromosome, and array density. The points correspond to either individual arrays or gaps (array length and gap length) or chromosomes (polycentromeric share and array density). The rearrangement rates are the rates along the terminal branches. **B**. Schematic representation of the models of fusion and fission rate on a species level, which revealed non-significant associations with polycentromere organisation variables. **C**. Scatterplots showing the relationship between the non-polycentromeric chromosome length and polycentromere share and density, with unique colours and trend lines for each species. **D**. The relationship between whether a chromosome is a fission product and the scaled median length of polycentromeric arrays on that chromosome, with glm fits for each species. **E**. Breakpoint regions are generally enriched for polycentromeres. The phylogeny was drawn using iTOL (https://itol.embl.de/)^40^.

Modelling rearrangement rate, the null model performed best, as assessed through leave-one-out cross-validation (Table S6). When modelling fusion rate separately, the model incorporating only array density had the best predictive performance (Bayesian stacking weight = 0.809, Table S6), though the posterior predictive intervals were broad (median width = 1.33 ⨯ 10^6^ compared to observed values up to 4.49 ⨯ 10^4^) indicating uncertainty in species-level predictions. Inspecting the summary statistics revealed that there was a negative relationship between scaled array density and fusion rate (mean: -0.530, 95% CI: -1.43 – 0.386) (Table S8). The other model with non-zero Bayesian stacking weight (0.191) was the amount model, incorporating both array density and polycentromere share. In this model, there was also a negative relationship between scaled polycentromere share and fusion rate (mean: -0.230, 95% CI: -1.14 – 0.711) (Table S8). The overlap of zero in these credible intervals and the broad posterior predictive intervals were unsurprising given the small number of observations (n = 30 species).

When modelling fission rate, the model with only the mean inter-array gap length had the best predictive performance (Bayesian stacking weight = 0.640). The summary statistics suggested a negative correlation between scaled inter-array gap length and fission rate (mean: -2.79, 95% CI: -6.17 – 0.531) (Table S9).The only other model with non-zero Bayesian stacking weight was the null model.

### 3.6 Chromosome size is a major determinant of candidate polycentromere organisation

We then asked whether the relationship between polycentromere distribution and chromosomal rearrangement may be patterned at the chromosome level. We first modelled chromosome size (Table S10, Table S11, Figure 3C). The model with only polycentromeric share of the chromosome as a predictor had the best predictive performance (Bayesian stacking weight = 0.715; median posterior predictive interval width = 1.84, relative to an observed range of 7.36 for scaled chromosome size). Larger chromosomes had greater polycentromeric share (mean: 0.127, 95% CI: 0.061 – 0.194). The model incorporating only inter-array gap length also had a non-zero Bayesian stacking weight; larger chromosomes have smaller gaps between polycentromeres (mean: -0.053, 95% CI: -0.096 – -0.011).

Larger chromosomes could be more likely to be a recent fusion product. When fitting models with the fusion and fission history of the chromosome as predictors, controlling for non-polycentromeric chromosome length, the only polycentromeric organisation variable with a credible interval not overlapping zero was median array length. Recently-split chromosomes had shorter candidate polycentromeres (mean: -0.118, 95% CI: -0.224 - -0.010) (Figure 3D, Table S12, Table S13). Chromosome length was significantly positively correlated with polycentromeric share and array density and significantly negatively correlated with inter-array gap length, demonstrating that the previous model comparison results are robust to controlling for fusion and fission history of a chromosome (Table S12).

### 3.7 Enrichment of inter-chromosomal rearrangement breakpoints in candidate polycentromere arrays

We wondered whether we could identify signatures of candidate polycentromeres having directly facilitated rearrangement events. By comparing genomes across species, we identified the regions of chromosomes that contained a fusion between ancestral chromosomes, or that were the site of a fission in closely related species. We found a significant enrichment of candidate polycentromeres in the fission regions of three of the twelve *Carex* species analysed, and a significant depletion was identified in two. Regarding fusion regions, a significant enrichment of candidate polycentromeres was identified in seven of the twelve *Carex* species (Figure 3E, Table S14). Enrichment of rearrangement breakpoint regions in coding sequences and TEs was also tested, but there were no consistent patterns (significant relationships in Table S15).

## 4. Discussion

### 4.1 Ancestral linkage group inference allows quantification of inter-chromosomal rearrangement rates from genomic data

Chromosome-level genome assemblies are essential for a comprehensive understanding of inter-chromosomal rearrangements. Here, we demonstrate this in the sedges and rushes, using 36 high-quality assemblies to reconstruct the ancestral linkage groups of a rapidly rearranging group and to accurately identify and quantify rearrangements along internal branches of our phylogeny. These analyses provide an important perspective on the extraordinary rearrangement rates in the cyperid clade. Quantifying rearrangement rates along internal branches using ALG reconstruction is expected to lead to some degree of underestimation, as syngraph cannot resolve complex rearrangement histories^42^. In the part of the tree with the greatest rearrangement rate, *Carex*, taxon sampling was also densest and branch lengths were shorter, minimising unobserved events.

Rearrangement rate estimates were notably high. These high rearrangement rate estimates, such as 1.31 inter-chromosomal rearrangements per million years in *Carex*, are some of the first estimates from genomic data. In other clades, rearrangement rates have been calculated using karyotypes and are lower than in *Carex*. For example, in birds, a Markov modelling approach estimated 0.12 fissions and 0.20 fusions per million years^78^. More similar to the high rearrangement rate of *Carex* is the holocentric butterfly genus *Erebia*, which is known to have exceptionally high rates of rearrangement among Lepidoptera with approximately 1.37 rearrangement events per lineage per million years^79^. Evidently, especially when considering the potential for rate underestimation, the rearrangement rate in *Carex*, and sedges overall, is remarkable.

The *Carex* species studied have chromosome numbers ranging from 22-54 (Figure 1), with most species having ∼30. Given the high rates of rearrangement in the clade this pattern could be due to either stabilising selection for an optimal karyotype, perhaps to maintain an optimal recombination rate across the genome^73^, or a general balance between fission and fusion. We found that a model with fusion probability fixed on each branch, at *p* ≈ 0.54, was sufficient to explain observed tip karyotype variance, and there was no clear tendency for fusion or fission probability to change based on the number of ALGs at the parent node.We found higher rearrangement rates in polycentric compared to monocentric clades, though our analysis is limited by the number of *Juncus* genome assemblies. There is extensive variation in rate within the holocentric groups. The polycentric Juncaceae genus *Luzula*, had a much lower rearrangement rate (0.182 rearrangements Myr^-1^ since the LCA of *Luzula* and *Juncus*) than did the clade of *Carex* and relatives. In the asatellitic holocentric Lepidoptera, rearrangement rate also varies greatly; most taxa retain an ancestral karyotype of n = 31, but some clades have experienced extensive rearrangement^80^. There seem to be factors other than lacking a monocentromere that promote rearrangement in certain clades. These could include population genetic effects or how the polycentromeres are organised. Future work could incorporate the comparison between clades exhibiting different degrees of karyotype stability in meiosis, once such a cytological dataset becomes available.

### 4.2 Polycentromeric drive may act on karyotype, centromere sequence, and centromere architecture

The rapid evolution of centromeres despite their important role inspired the centromeric drive hypothesis: alterations in satellite sequence or length could result in biased segregation into the gamete during asymmetric meiosis, providing a substrate for selection^18, 81^. Often the centromere with more or stronger microtubule attachments due to increased length or differing sequence is preferentially retained by the gamete^82^. However, if it is the pole in the polar body which has higher microtubule density, longer centromeres will be more likely to end up in the polar body, and drive could favour shorter centromeres^83^. Centromeric drive has been extended to accommodate holocentric species. In the absence of centromeric sequence features, “holokinetic drive” acts to increase or decrease the number of microtubule attachments simply through changing the length of the whole chromosome, through modulating repeat content, or through chromosomal rearrangements^19^. This theory was originally proposed to explain chromosomal evolution in sedges and rushes, before the polycentromeric nature of their spindle attachment was discovered^12^.

It has been noted that the satellite-based centromeres of polycentric species may allow centromeric drive to occur^84^. In accordance with this, we put forward a modification to existing drive models, which we refer to as “polycentromeric drive” (Figure 4). This combines aspects of both classical centromeric drive and holokinetic drive. We would expect evolution of polycentromeric satellite sequence to modulate the number and strength of spindle attachments. We could also expect changes in chromosome length through inter-chromosomal rearrangements, as fusions would increase the kinetochore extent on a chromosome, and fissions would decrease it. We might also expect concerted evolution in the density and length of polycentromere arrays. In species with an apparent cluster-like distribution of kinetochores, like *Luzula sylvatica* or *Rhynchospora pubera*^74, 85^, this may modulate total kinetochore extent and the number of microtubule attachments. In polycentric species where the kinetochore seems to assemble in a more continuous, line-like manner^11, 86^, density and array length may still affect kinetochore stability.

**Figure 4.**
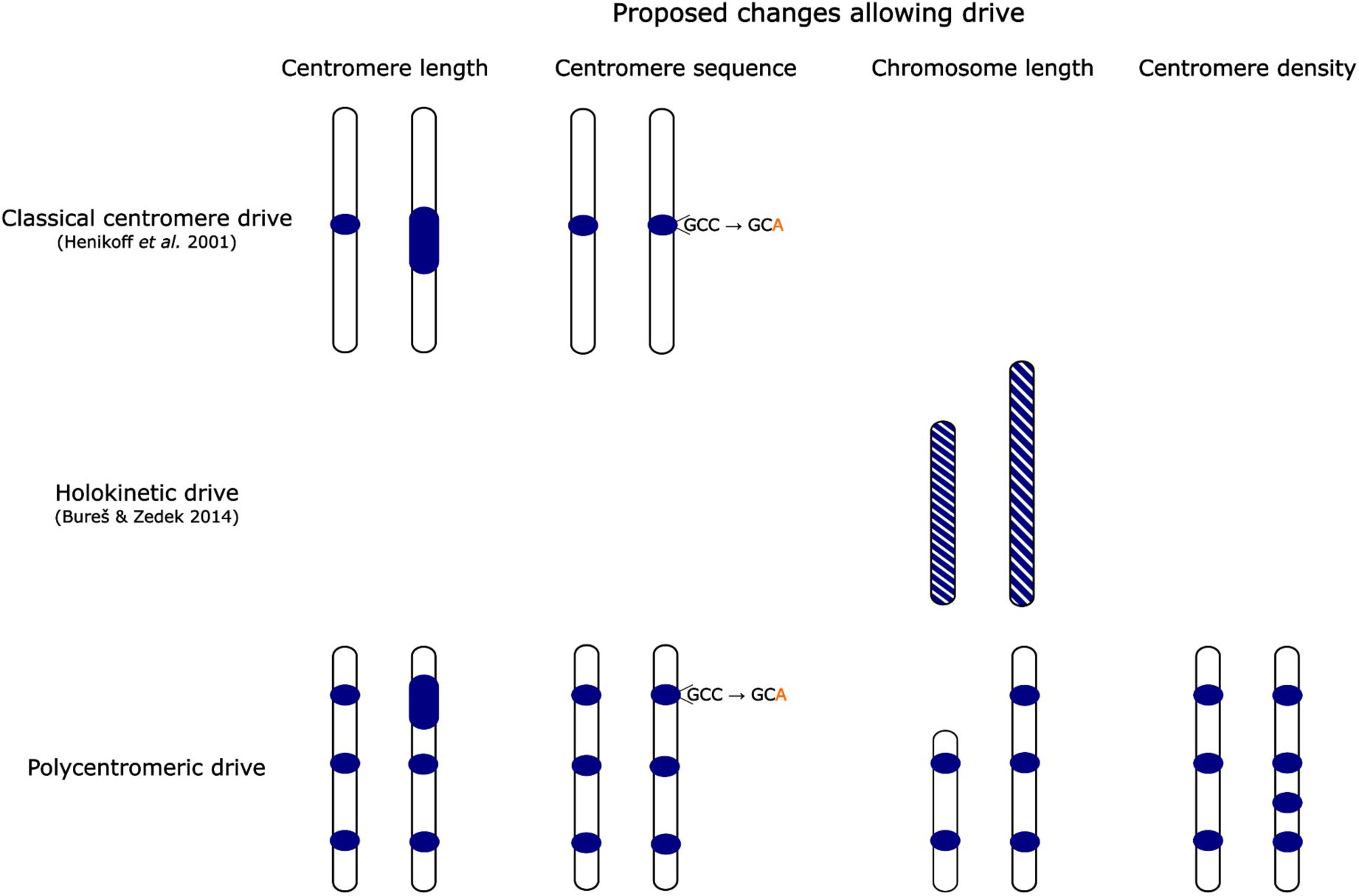
A comparison of classical centromere drive^18^, holokinetic drive^19^, and polycentromeric drive. Under classical centromere drive, changes in overall centromeric size or sequence can bias inheritance into the gamete in asymmetric meiosis. Under holokinetic drive, a change in the length of the chromosome will likely change the extent of the kinetochore, also potentially biasing inheritance into the gamete. Under polycentromeric drive, changes in individual centromere array size, sequence, density, and the length of the chromosome as a whole could all potentially bias inheritance.

Polycentromeric drive should be especially effective in sedges, as meiosis is asymmetric in both sexes^20^. Indeed, our annotation of candidate centromeric satellites revealed rapid sequence turnover, in line with predictions under polycentromeric drive. Although we found no evidence to suggest polycentricity has evolved in the Cyperaceae more than once, we were able to classify candidate polycentromeric satellite repeats of 29 sedge species into 18 families. However, it is notable that the same satellite families (*Carol* and *Sandra*) were found constituting polycentromeres across all *Carex* species studied. Persistence of sequence homology over tens of millions of years suggests occasional purifying selection acting on polycentromeres when the genome is not invaded by a new dominant satellite.

The high rates of inter-chromosomal rearrangement, modulating chromosome (and kinetochore) size and number^87^, may also be the product of polycentromeric drive. In concordance, previous studies in *Luzula* find that most species have either the ancestral 12 pairs of chromosomes or a derived 24, indicative of drive oscillating between favouring more/shorter kinetochores and fewer/longer kinetochores^87, 90^. However, drive-driven rearrangement is difficult to disentangle from the possibility of polycentromeric architecture directly facilitating rearrangement, as repetitive DNA is particularly vulnerable to the double-strand breaks (DSBs)^88, 89^ required for fission and fusion. Indeed, Zhang *et al.*^17^ found that polycentromeric *Tyba* arrays are hotspots for rearrangements in the genus *Rhynchospora*. We also found a general enrichment of polycentromeres in breakpoint regions across cyperids. High rearrangement rate may be a product of both polycentromeric drive and polycentromere-facilitated breakage.

We identified great variation in the architecture (i.e. length and density of the satellite arrays) of candidate polycentromeres. The mean length of the satellite arrays ranged from 8 kb (*Trichophorum cespitosum*) to 128 kb (*Schoenus nigricans*), while density ranged from 3 (*Cyperus fuscus*) to 38 (*Luzula pallescens*) arrays per 10 Mb. Such variation could be a signature of drive, but the exact mechanism involved likely depends on the dynamics of kinetochore assembly. In polycentric species in which kinetochores assemble in a discrete, clustered manner along the chromosomes, increasing length and density of the individual arrays could increase the number of spindle fibre attachments, biasing segregation. In support of this, kinetochore assembly in *Luzula sylvatica* is known to occur in a clustered pattern^74^, and we found that this species has especially high array density and total centromeric satellite sequence.

In comparison, the kinetochores of some other cyperid species assemble as a continuous line (often a groove) along the condensed chromosome^11, 12, 86^. In these cases, centromeric array length or density may not correlate with the extent of the kinetochore as directly^87^, but these variables may still be important for the number or strength of the spindle attachments. This is supported by our observation that larger chromosomes have greater total candidate centromeric sequence length and greater array density, even when these variables are normalised by chromosome length. This may be a response to a superlinear relationship between chromosome length and the mechanical demands of segregation in this clade, and suggests that array length and density do have a bearing on the strength of the kinetochore. Polycentromere architecture may thereby act as a substrate for polycentromeric drive in cyperids, though future work should aim to validate this experimentally by observing the segregation frequencies of homologous chromosomes with different polycentromere architectures.

Our results also suggest that the relationship between polycentromere architecture and chromosome size may in turn influence the stability of inter-chromosomal rearrangements. While longer chromosomes appear to have a greater share of their length occupied by centromeric sequence, when they split the fission products would end up with disproportionately much centromeric sequence. This disproportion may interfere with spindle fibre geometry and result in improper segregation. Indeed, we found that chromosomes that were recent fission products had shorter candidate polycentromere arrays, even when controlling for chromosome size, suggesting that they might be derived from chromosomes with shorter centromeres in the first place.

Too much centromeric sequence may also be problematic for larger cyperid chromosomes, affecting the stability of interchromosomal fusions. If more centromeric sequence means a greater number of spindle attachments, there may be a greater risk of at least some spindle attachments being misoriented, attaching to the opposite pole to the rest, resulting in chromosome tearing and missegregation. There is precedent for this in larger kinetochores being more likely to attach to both poles (merotely) in the monocentric Indian muntjac^91^. Consistent with this possibility is the trend of lower fusion rate in cyperid species in our dataset with greater candidate polycentromere share and density; fusions result in a larger kinetochore and perhaps more spindle attachments per chromosome, but the risk of merotely is lowered if the fusing chromosomes have less centromere sequence to begin with.

Polycentromere architecture also appears to influence the coding sequence and TE landscape of the chromosome. In most species of Cyperaceae in our dataset, we found that gene density decreased and TE density increased with proximity to candidate centromeric arrays, the same trend as is typically seen in monocentric species, and often explained by heterochromatin accumulation and recombination suppression at centromeres^76, 92–95^. In contrast, we found an enrichment in gene density and decay in TE density close to candidate polycentromeres in our two *Rhynchospora* species, which might suggest a history of polycentromeres and/or peripolycentromeric regions being recombination hotspots in this genus, even though extant populations of *R. tenuis* are achiasmatic^96^. This difference may be partly due to the comparatively high density of polycentromeres in *Rhynchospora* compared to other cyperid species. Selection might be acting to reduce the extent of pericentromeric heterochromatin associated with each polycentromere, so that too much of the chromosome is not heterochromatinised. This may then allow the relationship to be tipped in the opposite direction as the repetitive nature of polycentromeres results in a locally higher frequency of DSBs and DSB repair through homologous recombination, with a gene-rich and TE-poor signature. Nevertheless, correlating sequence features like TEs with polycentromere position cannot establish with certainty a causal direction; future work involving reconstruction of individual TE insertion histories could clarify this relationship.

Our investigation into the causes and consequences of the variation in polycentromeric architecture has two main limitations. Firstly, our annotation of polycentromere-related satellite arrays has not been validated experimentally in this study, though our method does resolve the same centromeric satellites as have been inferred for genera which have been analysed cytogenetically. Secondly, it is possible that measures of polycentromere architecture may depend on assembly quality. However, we expect this effect to be relatively minimal as individual polycentromere arrays tend to be shorter than monocentromeres, and can therefore be handled well by long read assembly. We anticipate that future refinement of polycentromere annotations through experimental validation and advances in assembly approaches for repetitive regions should be able to resolve the relationship between polycentromere architecture and genome evolution with greater resolution.

### 4.3 On the loss of polycentricity

Monocentricity is hypothesised to be the ancestral state for eukaryotes, and despite holocentrics (including polycentrics) accounting for ∼20% of species^8^, a reverse transition has never been documented^70^. The monocentric *Juncus* could represent a reversion from the polycentricity seen in other Juncaceae, but this cannot be distinguished from two independent origins of polycentricity in the cyperid clade given our phylogeny. Furthermore, we could identify neither orthologues of *Juncus* monocentromeric satellites in other cyperids, nor orthologues of polycentromeric satellites in *Juncus*, which is unsurprising given rapid satellite turnover in the clade.

However, our annotation of candidate centromeric satellites in *Carex myosuroides* did reveal arrays of the 177 bp satellite *Kelly* which resemble monocentromeres. These arrays are associated with decreases in gene density and increases in TE density in the pericentromeric regions. Furthermore, the constituent satellite repeats are of strikingly similar length to the main centromeric satellites in *Arabidopsis thaliana* (178 bp)^97^ and humans (171 bp)^98^, which may be relevant for nucleosome wrapping. Though the monocentric behaviour of the *Kelly*-harbouring chromosomes needs to be confirmed using molecular cytogenetics, a reversion to monocentricity would provide additional power for the study of the causes and consequences of changes in centromeric organisation. The same is true for the apparent loss of satellite-based centromeres altogether in *Cyperus rotundus*, an arguably even more dramatic shift, though the centromere architecture in this species also requires experimental verification.

## 5. Conclusions

Explaining the causes and consequences of derived traits is difficult when there are only a limited number of transitions between discrete states. This has been the case with holocentricity; while holocentric organisms represent ∼20% of eukaryotes^8^, the relatively low number of independent transitions from monocentricity to holocentricity makes comparative analyses weakly powered. Therefore, studying polycentric organisms which exhibit continuous variation in their centromere organisation provides important insights. The interplay between holocentricity and chromosomal evolution has implications for adaptation and speciation^7, 99^, but our study indicates that the relationship between centromere organisation and inter-chromosomal rearrangements is not straightforward. Though having multiple centromeres theoretically stabilises fission and fusion products, having more centromeres does not always mean higher rearrangement rates, as having too many also seems destabilising. This may be partly because polycentromere length scales superlinearly with chromosome size, and deviations from this relationship could interfere with spindle geometry.

The superlinear relationship between chromosome size and centromere size is itself unexpected given that studies in monocentric species reveal monocentromere size scales only linearly or sublinearly with chromosome size^100^. In other ways, though, our candidate polycentromeres behave like monocentromeres. For example, polycentromere units appear to be recombination coldspots given the signature of gene density reduction and transposable element enrichment closer to the centromeres, except for in the species with the densest centromeric organisation. Furthermore, drive still appears to be acting to promote centromeric satellite sequence turnover, as well as changes in polycentromere architecture and chromosome size. Additionally, transition between monocentricity and holocentricity may not be a one-way street, with the discovery of putative neo-monocentromeres in *Carex myosuroides*, a finding that would be good to validate experimentally. The cyperid clade has thousands of species, and we should expect continued chromosome-level genome assembly to reveal more variation in centromeres, lend more support to observed global patterns, and provide greater resolution for investigating subclade-specific relationships between centromeres and inter-chromosomal rearrangements.

## Acknowledgements

We would like to thank the many colleagues in the Tree of Life Programme at the Wellcome Sanger Institute, and the Henderson Lab at the University of Cambridge, for advice on methods, interpretation, and visualisation. This work would also not have been possible without those behind genome sequencing and assembly for the Darwin Tree of Life project, to whom we are also indebted. We are also grateful to Nathan Riley for advice on and for performing some Helixer annotations, and to André Marques for providing k-mer spectra for *Rhynchospora tenuis*, *R. breviuscula*, *R. pubera*, and *Juncus effusus*. We would also like to thank the three anonymous reviewers for their helpful comments.

This research was funded by Wellcome through core funding to the Wellcome Sanger Institute (206194, https://doi.org/10.35802/206194) and the Darwin Tree of Life Discretionary Award (218328, https://doi.org/10.35802/218328). For the purpose of Open Access, the author has applied a CC BY public copyright licence to any Author Accepted Manuscript version arising from this submission. The funders had no role in study design, data collection and analysis, decision to publish, or preparation of the manuscript.

## Data and code availability

Scripts used for analysis and visualisation, as well as input data, can be found at github.com/Obscuromics/cyperids.

## Author contributions

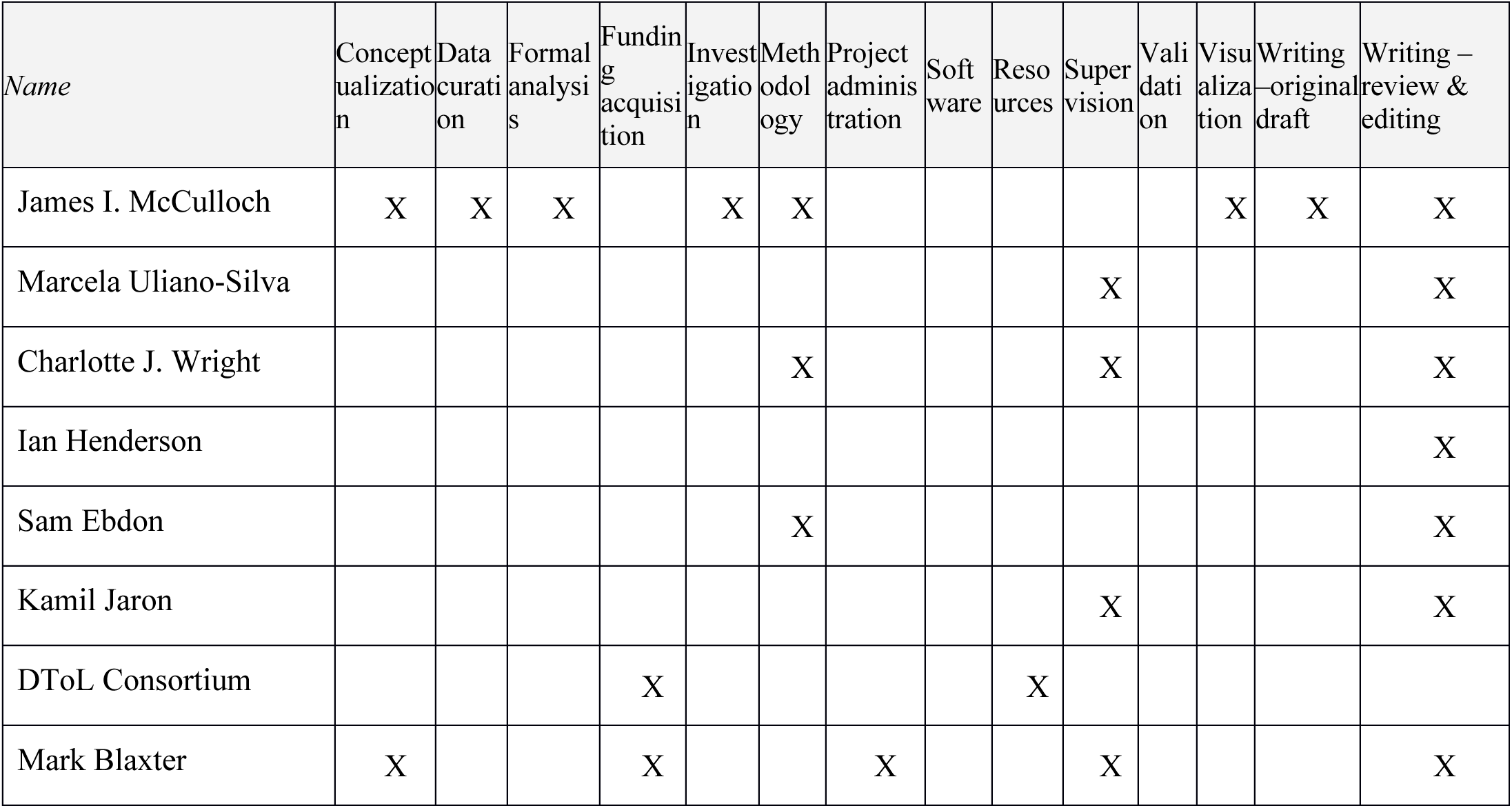

