## Supplementary Text for "Rapid chromosomal evolution and polycentromeric drive in sedges and rushes"

#### Supplementary Text 1. The dataset and phylogenetic reconstruction.

We downloaded chromosome-level genomes for 44 species in the families Cyperaceae and Juncaceae[21–29] (all species released on NCBI or ENA to November 2025). We included two additional Poales species as outgroups: *Oryza sativa* (Poaceae: Oryzoideae) and *Ananas comosus* (Bromeliaceae) (Table S1)[30, 31]. Single copy orthologues were identified using *BUSCO* (Benchmarking Universal Single Copy Orthologs) v 6.0.0, the poales_odb12 dataset ($n$ = 6282)[32] and MetaEuk[33] (Figure S1). As whole genome duplications, complicate ancestral linkage group inference, we removed eight assemblies from the dataset that had fewer than 75% complete and single-copy BUSCOs (Table S1). This resulted in a final dataset of 38 assemblies: 36 cyperids  plus two outgroups (Table S1).

Single-copy BUSCOs from the 38 taxa were converted to multisequence amino acid FASTAsusing *busco2fasta.py* (https://github.com/lstevens17/busco2fasta), and filtered to 80% taxonomic coverage, retaining 5596 orthologues. Each set of sequences was aligned using *MAFFT* v. 7.526[34],trimmed using *trimAl* v. 1.5.0 (-gt 0.8 -st 0.001 -resoverlap 0.75 -seqoverlap 80)[35]; (74 sequences removed). The remaining 5522 sequences were concatenated into a supermatrix using *geneStitcher.py* (https://github.com/ballesterus/Utensils). Both the concatenated supermatrix and the individual alignments were used to construct phylogenetic trees using *IQ-TREE* v. 2.4.0 (-m LG -B 1000)[36, 37]. Gene concordance factors (gCFs) were calculated using *IQ-TREE* to compare the phylogenies produced using the concatenated supermatrix and the individual orthologues[38]. Weighted ASTRAL[39] (*ASTER* package v1.23) was used to infer a summary tree from the individual gene trees. The resulting topologies were identical (Figure S2).

### Supplementary Text 2. The effect of modulating the -m parameter on the syngraph inference.

Changing the -m parameter modulates the minimum number of markers required to count a rearrangement, which can affect the number of ancestral linkage groups inferred. This value was tested between 1 and 100 and the distribution of markers in each inferred ALG was plotted at three nodes: the basal node of the cyperid clade (which includes Cyperaceae and Juncaceae), the base of the Cyperaceae, and the base of the Juncaceae (Figure S3). In general, increasing -m resulted in fewer markers being classified as part of an ancestral linkage group. In contrast, too low an -m parameter resulted in prediction of larger numbers of rearrangements and typically predicted just a single ancestral linkage group for the cyperids or one harbouring the majority of markers. This probably occurs when the upper limit of the inferrable number of rearrangements is reached (see Mackintosh *et al.*, 2023 Fig. 2). We required a minimum of 33 markers to affect a rearrangement as this is the lowest value of this parameter which provides an inference not involving an ALG comprising >50% of the allocated markers in either the basal Cyperaceae node or Juncaceae nodes (Figure S3). Though inferences using higher -m parameter values also infer ALGs of which none harbour >50% of the allocated markers, they have fewer markers allocated to ALGs in total.

### Supplementary Text 3. Four models of fission and fusion probability.

***(i) "Fixed fusion probability" model***

The first model tests whether fusions and fissions occur with the same probability on all branches in the clade, and whether this is sufficient to explain observed variation in tip karyotypes. In this model, rearrangements were simulated as Bernoulli trials with fixed probability of fusion ($p$) calculated from observed fusion proportions. The number of fusions per branch ($F_i$) followed a binomial distribution dependent on the number of rearrangements per branch ($N_i$) and $p$:

$$F_i \sim Binomial(N_i, p), \qquad p = 0.54079254079$$

To assess the impact of the underestimation of rearrangement counts, the simulations were also run with the number of steps per branch multiplied by factors of 1.5 and 2. In addition, assuming that the observed distribution of tip chromosome number could be explained by a fixed fusion probability $p$ and underestimation of rearrangements, maximum likelihood estimation was used to estimate the value for the multiplier $m$. For values of $m$ from 0.1 to 10 (explored using continuous one-dimensional optimisation), 2,000 simulations of karyotype evolution were performed to produce a distribution of the log of variance in tip chromosome number. These distributions were approximated by a Normal distribution (hence the log-transformation of variance, on account of the right-skewed tip chromosome number variance distributions), and the log of the observed tip variance was used to find the maximum likelihood estimate for $m$.

$$\ell_{log}(m) = -\frac{1}{2}log(2\pi\sigma^2_{log}(m)) - \frac{(logV_{obs}-\mu_{log}(m))^2}{2\sigma^2_{log}(m)}$$

A full simulation with 10,000 trials was then run with the number of steps per branch multiplied by the maximum likelihood estimate for $m$.

***(ii) "Branch-heterogeneous" model***

An alternative model does not keep the fusion probability (*p)* equal for every branch, thereby testing whether allowing fusion probability to vary based on independent draws from a distribution of probabilities can replicate observed variation in tip karyotypes. In this model, branch-specific fusion probability ($p_i$) was drawn from a Beta distribution:

$$F_i \mid p_i \sim Binomial(N_i, p_i), \quad p_i \sim Beta(\alpha, \beta)$$

The hyperparameters were estimated using the observed number of rearrangements and fusion proportions by maximising the marginal likelihood under a Beta-Binomial model, which integrates over latent $p_i$. This naturally downweights observed fusion proportions from branches with low numbers of rearrangements, reducing noise and preventing fitting a biologically-unrealistic U-shaped Beta distribution. This would otherwise happen given that fusion proportions of 0 or 1 were commonly observed on branches with very low N.

$$\ell(\alpha, \beta) = \sum_i log\frac{B(F_i + \alpha, N_i - F_i + \beta)}{B(\alpha,\beta)}$$

$$\alpha = 5.16885..., \beta = 4.12471...$$

Karyotype evolution was simulated under the "fixed fusion probability" and "branch-heterogeneous" models as random walks. The random walks start at the base of the *Carex* phylogeny with 26 chromosomes (the number of *Carex* ALGs) and take $N_i$ steps per branch, the syngraph-inferred number of rearrangements. At each step chromosome number either decreases with probability $p_i$ (probability of fusion) or increases with probability (1 - $p_i$) (probability of fission). 10,000 simulations were performed for each model and variance in tip chromosome number calculated for each trial. The resultant distributions were compared to the observed tip karyotype variance.

***(iii) "Stabilising bias" model***

The third model tests for a force of "stabilising bias" which modifies fusion probability in order to maintain an optimum number of chromosomes. Per-branch fusion probability varied depending on the number of chromosomes at the parent node ($C_i$) compared to the "optimum" number of chromosomes ($C^*$), chosen to be 26, the number of *Carex* ALGs. As a sensitivity analysis, the analysis was also conducted with $C^* = 31$, the median observed tip karyotype. The baseline fusion probability ($p_0$), 0.54 in accordance with the "fixed fusion probability" model above, is modified for each branch in the direction that increases the probability of chromosome number changing in the direction of $C^*$. In other words, when $C_i$

> $C^*$ fusion probability increases and vice versa. The strength of modification is governed by the tuning parameter $\gamma$.

$$p_i(C_i) = logit^{-1}(logit(p_0) + \gamma(C^* - C_i))$$

The tuning parameter $\gamma$ was estimated with likelihood using the observed per-branch fusion frequencies ($F_i$) and the number of ALGs at internal nodes ($C_i$):

$$\ell(\gamma) = \sum_i [F_i \, log \, p_i(C_i) + (N_i - F_i) \, log \, (1 - p_i(C_i))]$$

***(iv) "Finite-memory heterogeneity" model***

This model tests whether there is autocorrelation in fusion probability along lineages, which could shape the distribution of chromosome numbers at the tips. Accordingly, the per-branch fusion probability $p_i$ is dependent both on the baseline fusion probability $p_0$ and the fusion probability along the parent branch, $p_{parent}$. The model also incorporates the parameter ρ, with larger values of ρ increasing the extent to which $p_i$ is influenced by $p_{parent}$, and the error term $\varepsilon_i$, adding some random noise.

$$\eta_i = (1 - \rho)\eta_0 + \rho\eta_{parent(i)} + \epsilon_i,$$

$$\text{where } \eta = logit(p) \text{ and } \epsilon_i \sim \mathcal{N}(0, \sigma^2)$$

As the probability model for $\eta_i$ is Normal, ρ and σ were estimated using the following log likelihood formula and observed fusion proportions:

$$\ell(\rho, \sigma) = \sum_{i=1}^{n} [-\frac{1}{2} log(2\pi\sigma^2) - \frac{(\eta_i - [(1-\rho)\eta_0 + \rho\eta_{parent(i)}])^2}{2\sigma^2}]$$

### Supplementary Text 4. Parsing the output of *CAP* to identify candidate polycentromeres

The putative satellite repeat which was the predicted polycentromeric repeat in the greatest number of species had a consensus length of 81 bp and was named *Carol* (***Car**ex* **polycentromer**e). This is predicted to be the main polycentromeric repeat in the larger of the two subclades which diverge from the *Carex* basal node in our phylogeny (Fig 1). It occurs on all chromosomes of all species in this subclade.

For all species in the large subclade, *Carol* also has an average CAP score (related to centromere probability) of >70, except for *Carex acutiformis*, *C. hirta*, and *C. rostrata*. Alongside *Carol*, *C. acutiformis* also has long arrays of a satellite with an 89 bp consensus sequence, found on most of the 39 chromosomes. Their presence is the probable cause of the lower average centromere probability of *Carol*; this satellite is usually found at the centromere ends. In *C. hirta*, despite *Carol* being present on all chromosomes, the satellite with the greatest average CAP score has a 143 bp consensus sequence, and is present on 26 of the 54 chromosomes, usually as telomeric arrays. An orthologous satellite is present in *C. rostrata*, with a consensus sequence of 144 bp, and is found in arrays at one telomere in each of 7 of the 35 chromosomes. *C. riparia*, which is the sister species to *C. rostrata* in our phylogeny and forms a triplet clade with *C. hirta*, has an average probability of 82 for *Carol* as the centromeric satellite, but also has arrays of the orthologous 144 bp satellite at one or both telomeres of 35 of the 36 chromosomes.

In the other main subclade of *Carex*, *Carol* does not appear to be the (only) main polycentromeric satellite. In the sister species *Carex myosuroides* and *C. littledalei*, *Carol* is also present, but it is absent from 14 of the 29 chromosomes of *C. littledalei* and 11 of the 29 chromosomes of *C. myosuroides*. *Carol2*is the satellite with the greatest centromere probability on 12 of the 16 chromosomes on which it is present in *C. littledalei*, but the satellite with the greatest average centromere probability has a 60 bp consensus sequence, here called *Marcus*. However, this is usually found on the same chromosome as *Carol* and is only found on three of the 14 chromosomes from which *Carol* is absent. Of the other 11 chromosomes from which *Carol2* is absent, seven have a satellite with a 125 bp consensus sequence having the greatest centromere probability, here called *Jerusha*. Of the remaining four, one has *Jerusha* present but a 105 bp satellite as the most probable centromeric satellite, one has *Jerusha* present but a 78 bp satellite as the most probable centromeric satellite, and one has *Jerusha* present but a 438 bp satellite as the most probable centromeric satellite. The only chromosome which lacks *Carol*, *Marcus*, and *Jerusha* has three types of repeat, in similar copy number, with 203, 128, and 438 bases. The one with 438 bases, here called *Mateo*, is considered most likely to be an polycentromere-associated repeat because it is regularly found on other chromosomes and is the most probable centromeric satellite in the aforementioned *Jerusha*-harbouring chromosome. In summary, there are four main putative types of polycentromere in *Carex littledalei*: *Carol*, *Marcus*, *Jerusha*, and *Mateo*.

In the *C. myosuroides* genome, *Carol* is the satellite with the greatest centromere probability on only two of the 18 chromosomes on which it is present. 13 of the 29 chromosomes have a

7 bp satellite, 5'-GTTTAGG-3' having the greatest centromere probability, including five of the 11 chromosomes from which *Carol2* is absent. This is typically a telomeric satellite variant, and in other cyperid species it occurs uniquely at the telomere, but in *Carex myosuroides* it has invaded the chromosomes away from the telomeres. It is found at regular intervals along most chromosomes, a pattern not seen in other genomes, supporting that this is not a signature of recent fusions. Further, the positions of these satellite arrays do not align with fusion points as assessed by synteny. Seven of the 29 chromosomes have a 224 bp satellite having the greatest centromere probability; however, it is localised to the chromosome ends, and is not found on chromosomes without *Carol* and/or 5'-GTTTAGG-3', and therefore may not have centromeric function. Five of the 29 chromosomes have a 177 bp satellite having the greatest centromere probability; it is present only in single arrays (on these five, as well as three further chromosomes where either *Carol* or 5'-GTTTAGG-3' have the greatest centromere probability) and therefore represents a putative neo-monocentromere, named *Kelly*. One of the remaining chromosomes has *Marcus* having the greatest centromere probability; it is also found on five other chromosomes. The final chromosome has a 440 bp sequence having the highest centromere probability; this sequence is orthologous to the 438 bp *Mateo* satellite of *C. littledalei*. In summary, the main putative centromeric satellites in *Carex myosuroides* are four polycentromeric repeats (*Carol*, 5'-GTTTAGG-3', *Marcus*, and *Mateo*) and one neo-monocentromeric repeat (*Kelly*). *Kelly* also occurs in *C. littledalei*, but it is only present as one short array (20-40 repeats) on each of four chromosomes.

The final four species of *Carex*, *C. divulsa*, *C. spicata*, *C. arenaria*, and *C. echinata*, are the sister clade to *C. littledalei* and *C. myosuroides* in our phylogeny but do not have *Carol* has a putative main polycentromeric repeat. Instead, the putative polycentromeric satellite is a repeat with a 124 bp consensus here named *Sandra*, present on all chromosomes of all four species, and with an average CAP centromere probability of >70 in all cases.

A much shorter variant of *Sandra* (30-31 bp), which seems to be the ancestral component of the secondarily homogenised 124 bp tetramer following divergence of the multiplicated monomer, is also one of the candidate polycentromeric satellites in *Eriophorum angustifolium*, *E. vaginatum*, and *Scirpus sylvaticus*, with a consensus sequence length of 30-31 bp. In both *E. vaginatum* and *E. angustifolium Sandra* is found on every chromosome, though both species also have a long satellite with a consensus sequence of 703 and 704 bp respectively, called here *Janet*, which has a high centromere probability on several chromosomes. This is especially the case for *E. angustifolium*, in which it has the highest average centromere probability. *Janet* is less common in *E. vaginatum*, but is joined by a 132 bp satellite, here called *Glenn*. Other satellite types have high centromere probability on certain chromosomes, but these appear to be localised expansions, except for a 12 bp satellite on *E. angustifolium*. This satellite typically forms arrays on one or both ends of the chromosome, and so may have telomeric rather than centromeric function. In *Scirpus sylvaticus*, *Sandra* is the most probable centromeric satellite on 15 of the 31 chromosomes, and a 40 bp variant of *Glenn* is the most probable centromeric satellite on a further seven. The other chromosomes have their most probable centromeric satellite being seemingly a localised satellite expansion, which always coexists with either or both of *Sandra* and *Glenn*.

*Sandra* is completely absent from *Trichophorum cespitosum*, which is the sister taxon to *Eriophorum* and *Scirpus*. Instead, *Glenn* appears to be the most dominant putative polycentromeric satellite. It has the highest centromere probability on 20 of the 52 chromosomes, and is present on all but three. Following that is a 123 bp satellite, which has the highest centromere probability and is found on all three of the chromosomes missing *Glenn*. This satellite has no orthologues in *Eriophorum* or *Scirpus*, and is named *Harmonica*. A 105 bp satellite has the highest centromere probability on six chromosomes and is named *Parker*. Finally, a 131 bp satellite has the highest centromere probability on a further six chromosomes and is named *BoDerek*. The other chromosomes appear to have their highest-probability satellite belonging to relatively localised expansions, such as a 22 bp repeat which forms arrays at one end of four chromosomes.

In *Cyperus fuscus*, a 325 bp satellite hereon called *Sayid* is the highest-probability centromeric satellite. In *Cyperus rotundus*, *Sayid* is not present, and most chromosomes have only the telomere satellite variant 5'-GTTTAGG-3', not invading away from the telomeres as it does in *Carex myosuroides*.

*Bolboschoenus planiculmis* and *x Bolboschoenoplectus mariqueter* have near-identical satellite landscapes. A 182 bp repeat, hereon named *Garrett*, is the satellite with the greatest centromere probability on 43 and 40 chromosomes respectively out of 54. In all other cases the highest centromere probability belongs to apparently localised repeat arrays or relatively long expansions of a 48 bp repeat on a minority of chromosomes, except for one chromosome of *x Bolboschoenoplectus mariqueter* on which the satellite with the greatest centromere probability is a different 48 bp satellite, hereon named *Colleen*. This repeat is consistently present in low frequency, occupying 38 of the chromosomes of *x Bolboschoenoplectus mariqueter* and 45 of the chromosomes of *Bolboschoenus planiculmis*.

In *Rhynchospora breviuscula* and *R. tenuis*, the satellite repeats with high centromeric probability have consensus sequences of 172 and 173 bp respectively, orthologous to the *Tyba* repeats previously characterised in this genus[12].

All 21 chromosomes of *Schoenus nigricans* have a 188 bp repeat, here named *Justine*, having the highest centromere probability. These repeats are present in relatively long arrays. 19 of these chromosomes also have a 168 bp repeat, here called *Sikowicz*, flanking at least one of these *Justine* arrays.

The results from CAP for the Juncaceae genomes mostly align with the centromeric repeats previously characterised in the literature[75]. *Juncus effusus* has centromeric regions harbouring *JefSAT1*, *JefSAT2, and JefSAT3*, which are grouped into the families *Amelia*, *Simms*, and *Cheyenne*, as all three are also found in *J. inflexus*. *J. squarrosus* has a distinct satellite with a 157 bp consensus sequence, named *Myrtle*. Both *Lusy1* (125 bp) and *Lusy2* (175 bp)[74] can be identified as polycentromeric-associated satellites in *Luzula sylvatica*, classified here in the families *Isaac* and *Jerry*, and are also found in *L. pallescens. L. pallescens* has a third satellite type with high centromeric probability, 104 bp, named here *Tony*.
