## Supplementary Figures for "Rapid chromosomal evolution and polycentromeric drive in sedges and rushes"

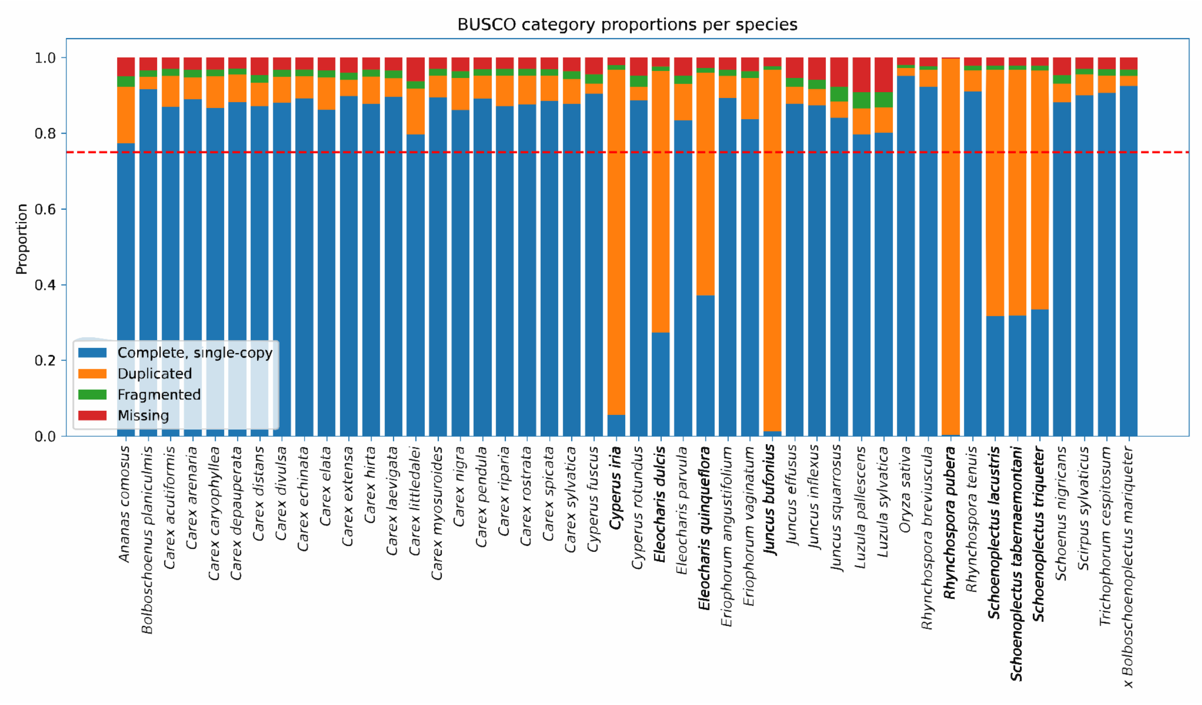


#### **Figure S1**. BUSCO metrics for assemblies used

For the initial 46 assemblies, the proportion of BUSCOs belonging to each category. The horizontal red line shows the threshold for inclusion: 75% of BUSCOs complete and single-copy.

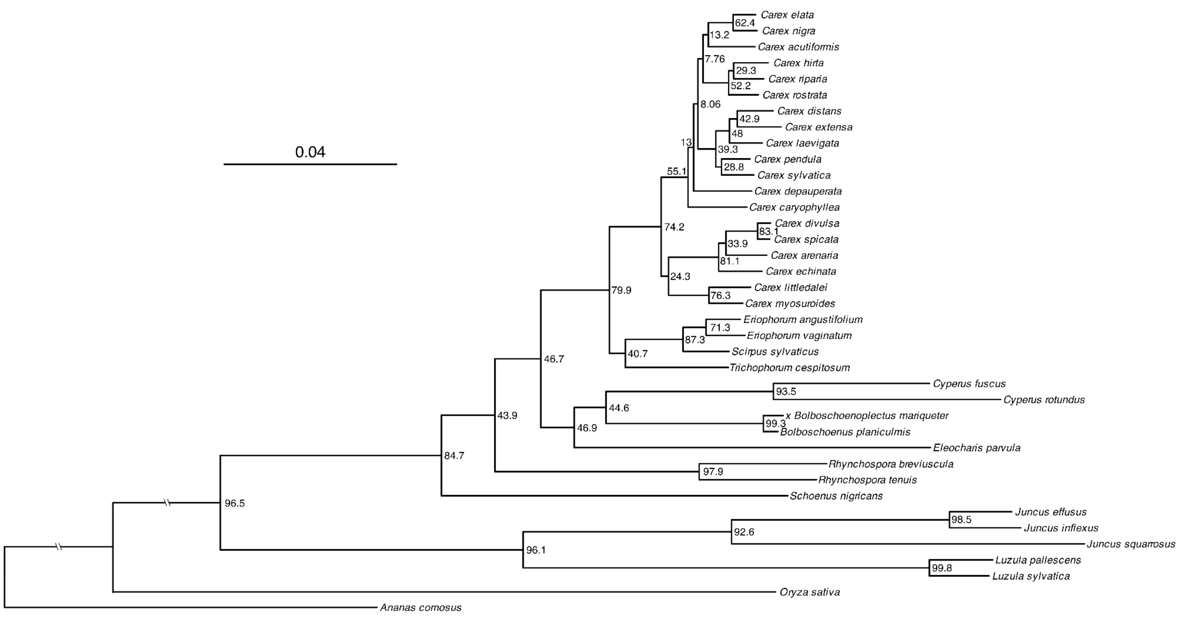


**Figure S2**. The phylogenetic tree used for the analyses, generated from a concatenated supermatrix of 5522 amino acid BUSCO sequences using *IQ-TREE*.

Node labels are the gene concordance factors (gCF), representing the percentage of decisive gene trees that support that node. All bootstrap scores are 100%. Despite low gCF values for some nodes, the weighted ASTRAL summary tree of the gene trees does not differ in topology. The scale unit is substitutions per site. Tree plotted using iTOL (https://itol.embl.de/)[40].

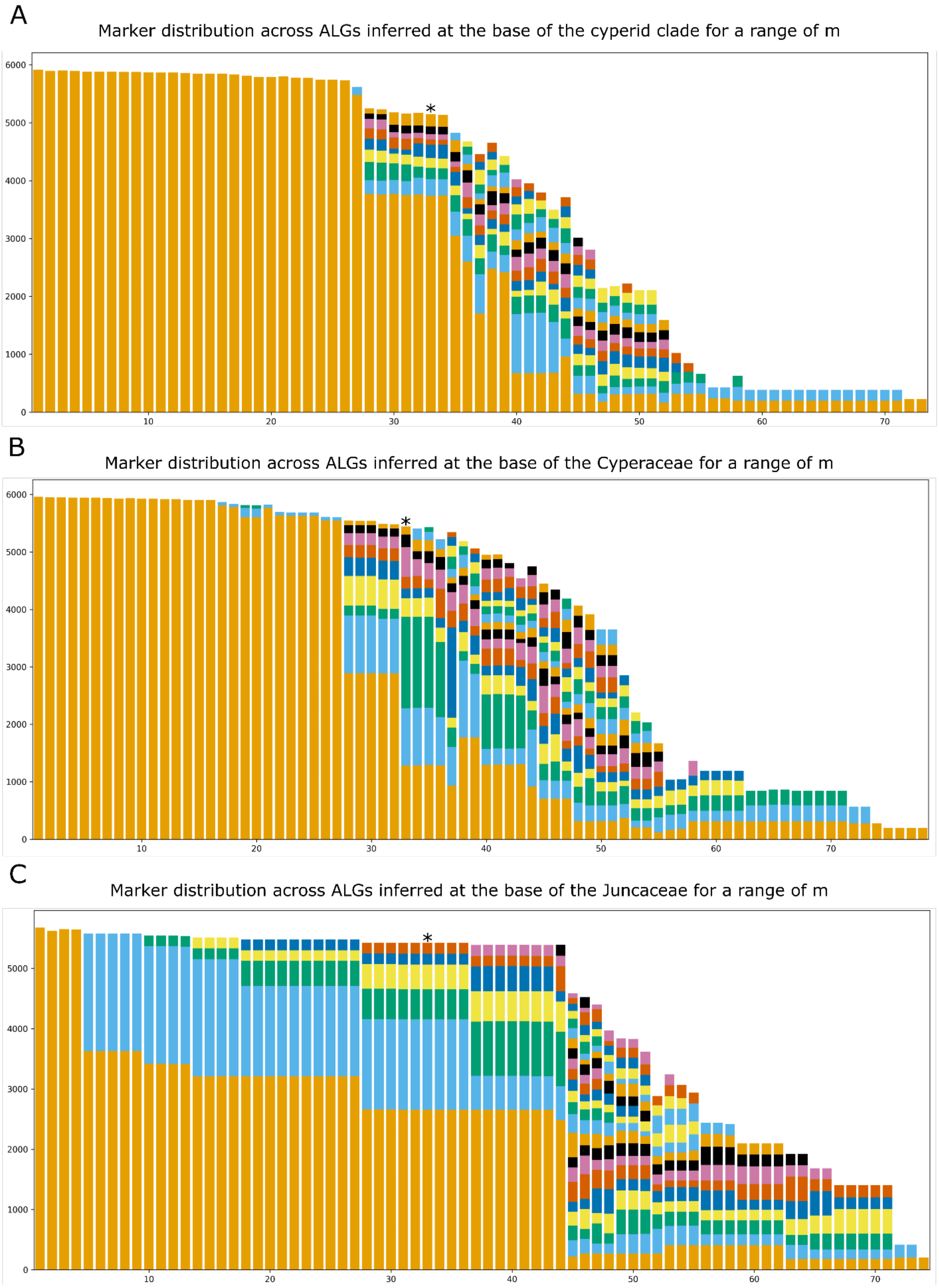


**Figure S3**. ALG marker assignment at ancestral nodes

Changing the -m parameter modulates the minimum number of markers required to count a rearrangement, which can affect the number of ancestral linkage groups inferred. The value of

-m was tested between 1 and 100 and the distribution of markers in each inferred ALG was plotted at three nodes: the basal node of the cyperid clade (which includes Cyperaceae and Juncaceae, panel **A**), the base of the Cyperaceae (**B**), and the base of the Juncaceae (**C**). In general, increasing -m resulted in fewer markers being classified as part of an ancestral linkage group. In contrast, too low an -m parameter resulted in prediction of larger numbers of rearrangements and typically predicted just a single ancestral linkage group for the cyperids or one ALG harbouring the majority of markers. This ancestral karyotype is likely predicted when the upper limit of the inferrable number of rearrangements is reached (see Mackintosh *et al.*, 2023 Fig 2.).

The value of m chosen for the downstream analysis, -m 33, is asterisked in each plot. This was chosen because for the Cyperaceae this is the lowest value which provides an inference not including one ALG comprising >50% of the markers.

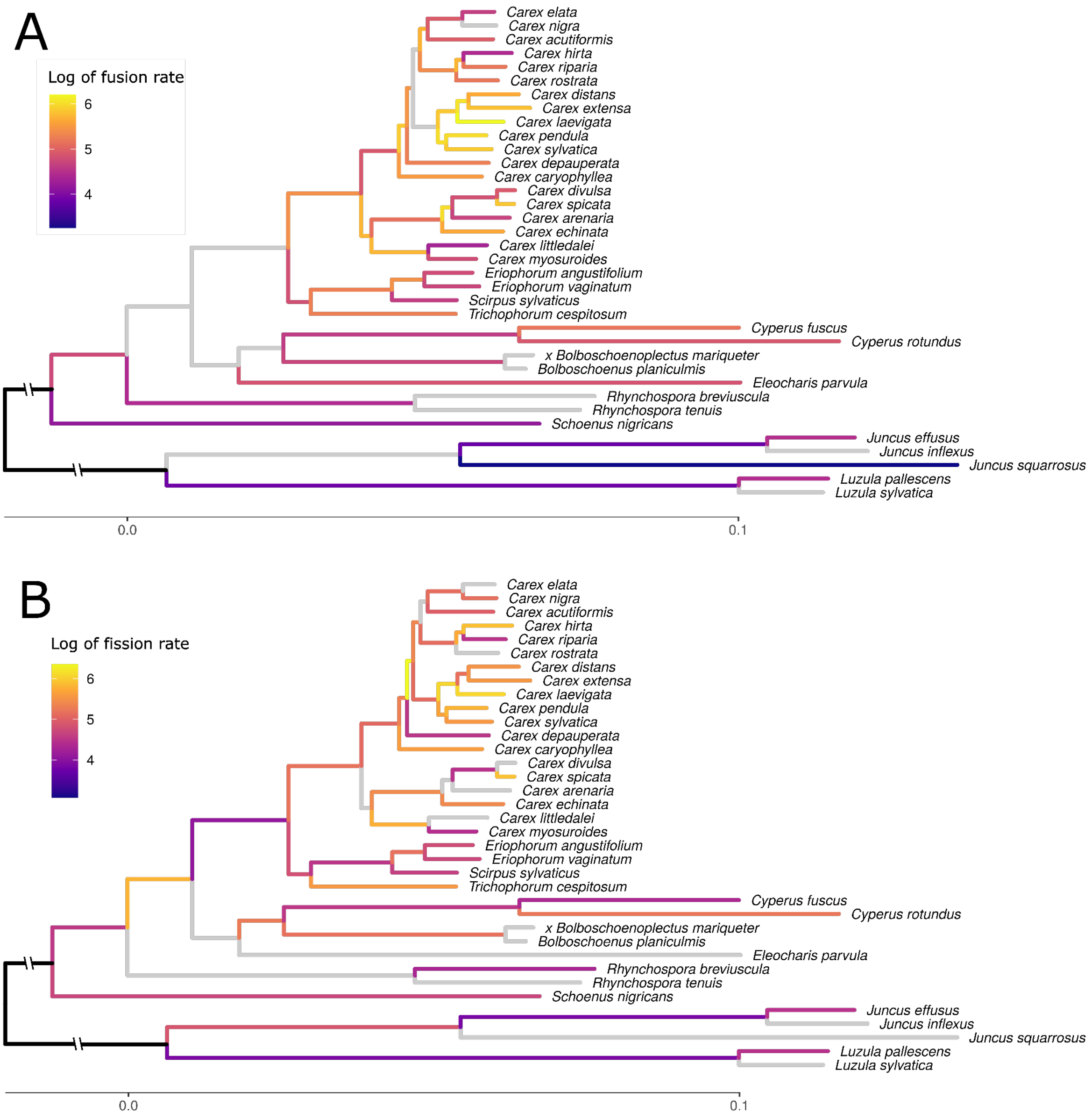


**Figure S4**. Rates of fusion and fission on the cyperid phylogeny.

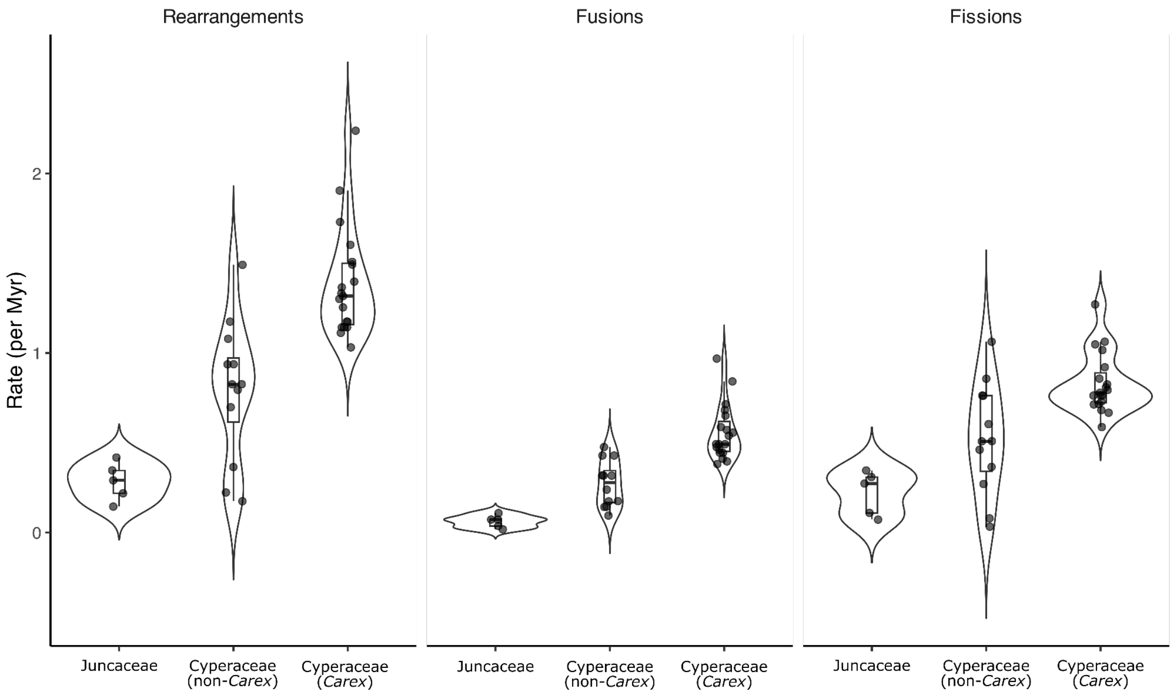


**Figure S5**. Violin plots showing the distribution of lineage-average rearrangement, fusion, and fission rates for Juncaceae and Cyperaceae, with Cyperaceae divided into non-*Carex* and *Carex* species.

For both non-*Carex* and *Carex*, the rate is calculated using the time elapsed since the base of the Cyperaceae in our tree. For all rates, the time elapsed is the median time indexed by TimeTree 5.

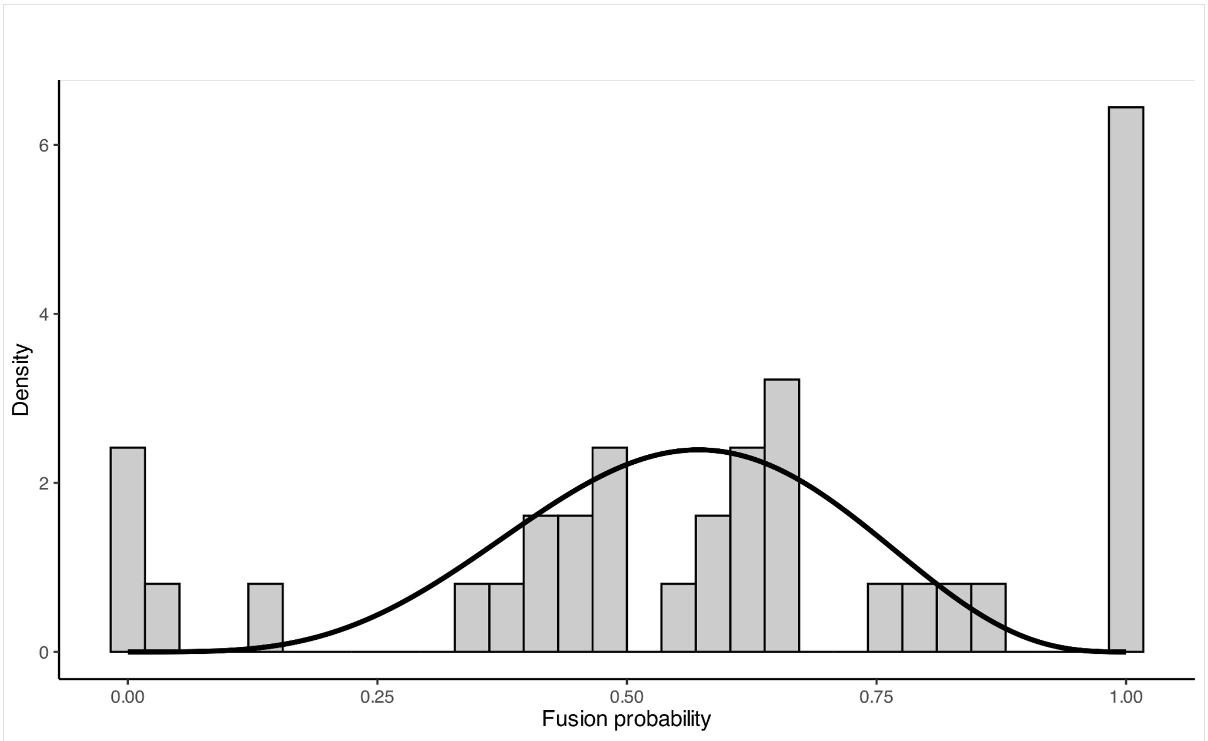


#### **Figure S6**. Per branch fusion probabilities in *Carex*.

A histogram of observed per-branch fusion probability (grey bars) in the *Carex* clade overlaid with the Beta distribution used for the branch-heterogeneous model of karyotype evolution, fitted by maximising marginal likelihood under a Beta-Binomial model.

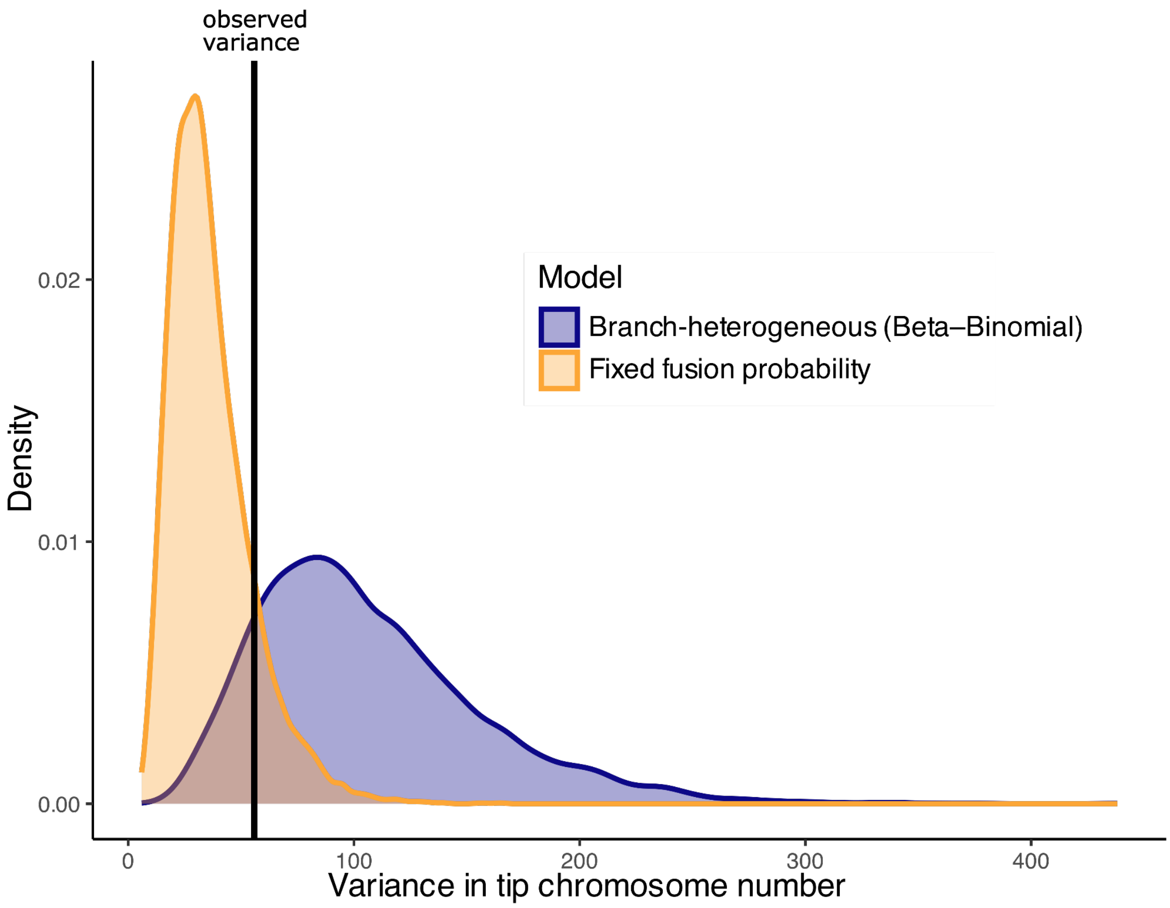


**Figure S7**. Comparing distributions of variance in tip chromosome number under the "fixed fusion probability" model and the "branch-heterogeneous" model.

Distributions of the variance in tip chromosome number from 10,000-trial simulations under two models: fusion probability fixed at 0.54 for each branch, and the "branch-heterogeneous" model, whereby fusion probability for each branch is drawn from a Beta distribution fitted using observed fusion probabilities. Both models have similar densities at the line denoting the observed variance in tip *Carex* karyotypes.

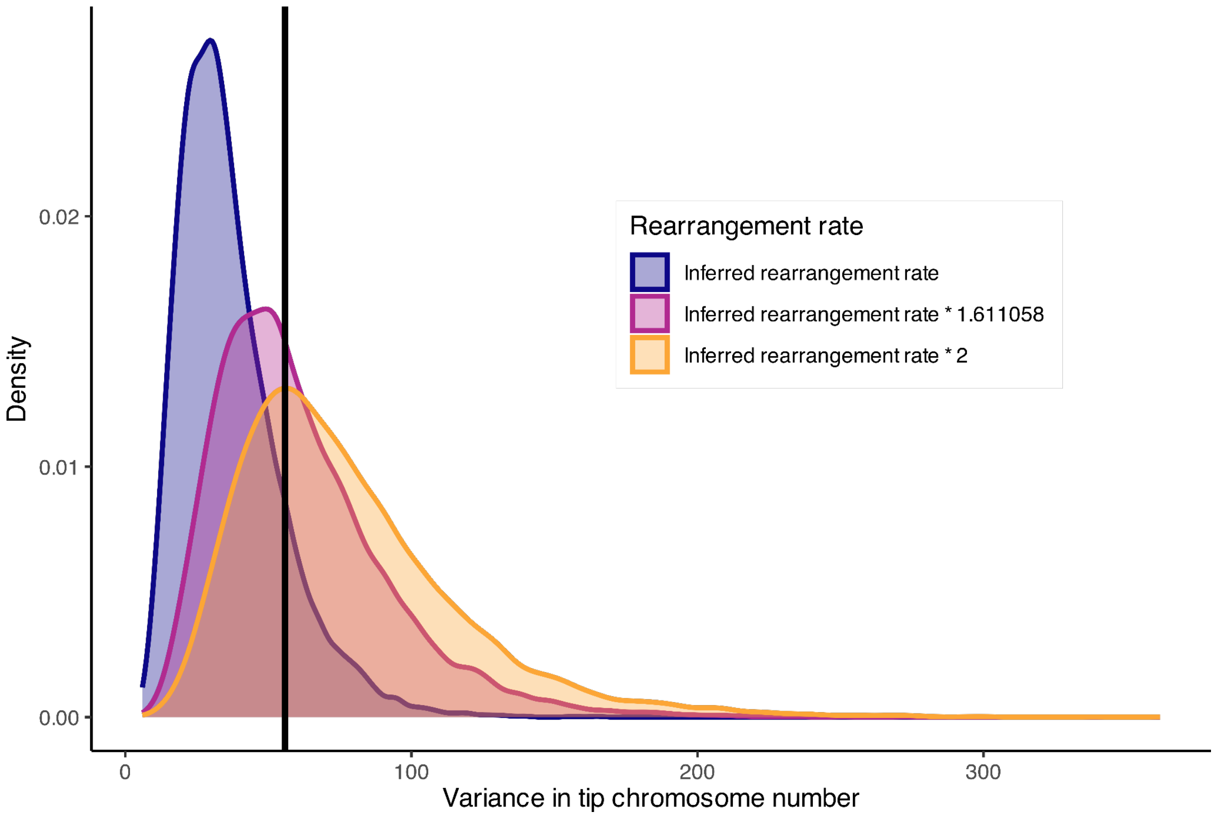


**Figure S8**. Distributions of the variance in tip chromosome number in *Carex* under models with different rearrangement rates.

Distributions of the variance in tip chromosome number from 10,000-trial simulations under three models, all with fixed fusion probability, but with the number of rearrangements along each branch either identical to that inferred by ALG inference, multiplied by a factor of 1.611058 (the maximum likelihood estimate for the rearrangement rate multiplier), or by a factor of 2. The observed variance in tip chromosome number is represented by the black line.

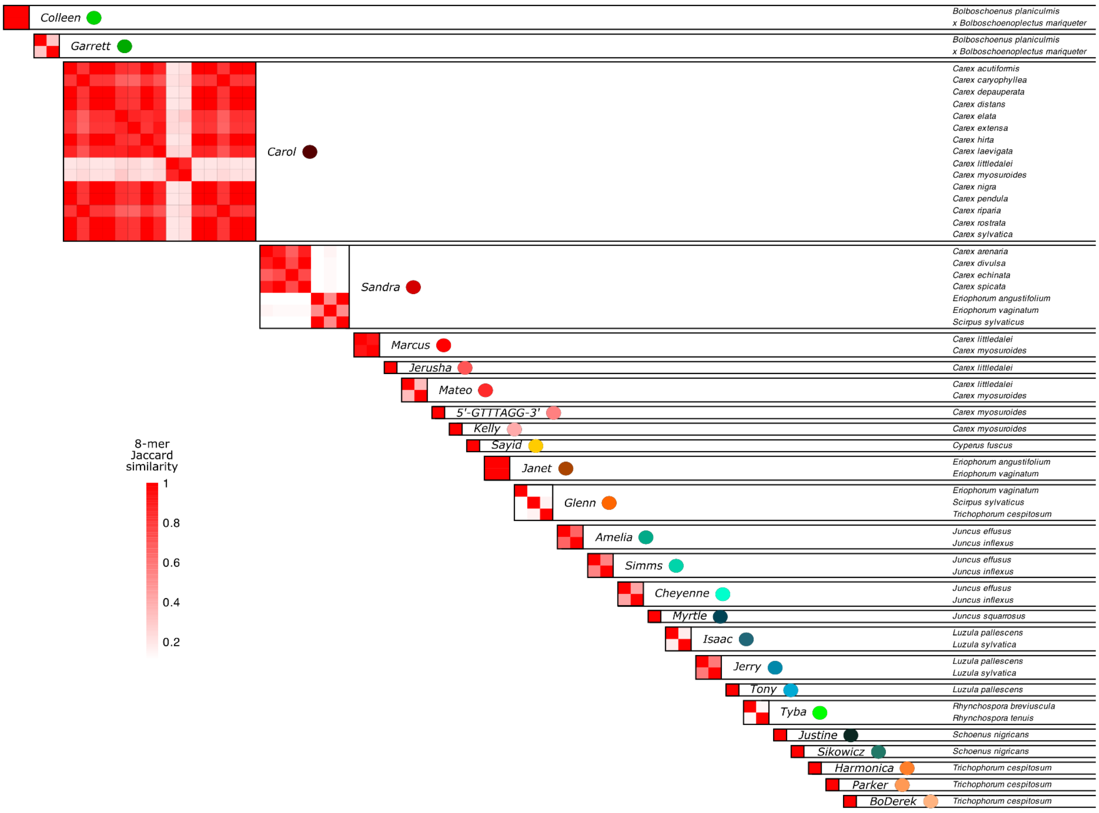


**Figure S9**. The similarity matrix from Figure 2, with species labels.

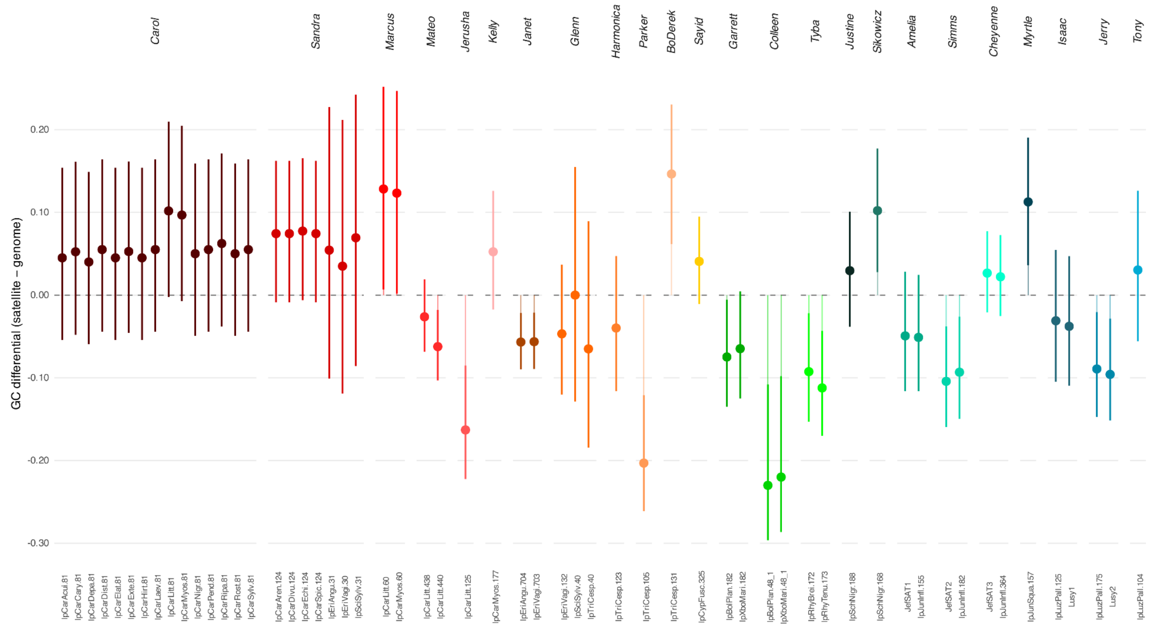


#### **Figure S10**. GC differential for each putative centromeric satellite.

We calculated GC differential by subtracting each species' reference genome GC content from the GC content of the satellite consensus sequence.

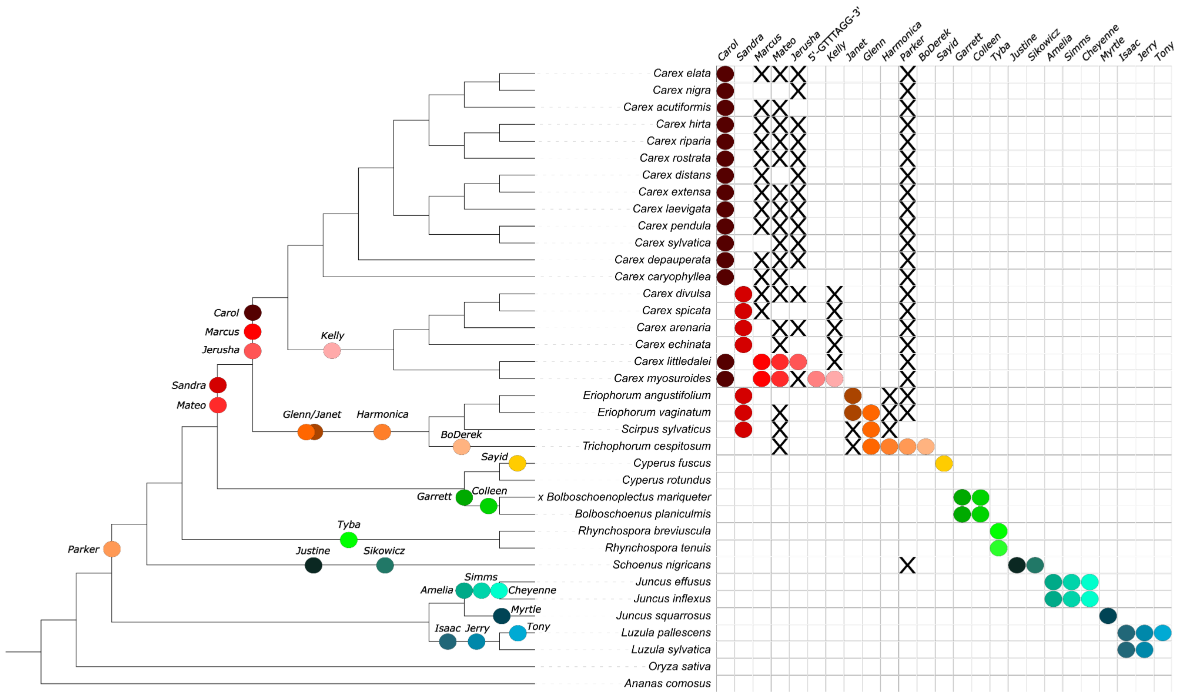


### **Figure S11**. Parsimonious origins of polycentromeric satellites.

We performed a BLAST of all satellite repeats identified by TRASH2 against the candidate polycentromeric satellite consensus sequences from each species. Hits with an E-value of at least 1e-3 and a coverage of 40% of the shorter sequence are denoted by an X. The most parsimonious origin of each satellite given these hits is annotated on the phylogeny. The phylogeny was made using iTOL (https://itol.embl.de/)[40].

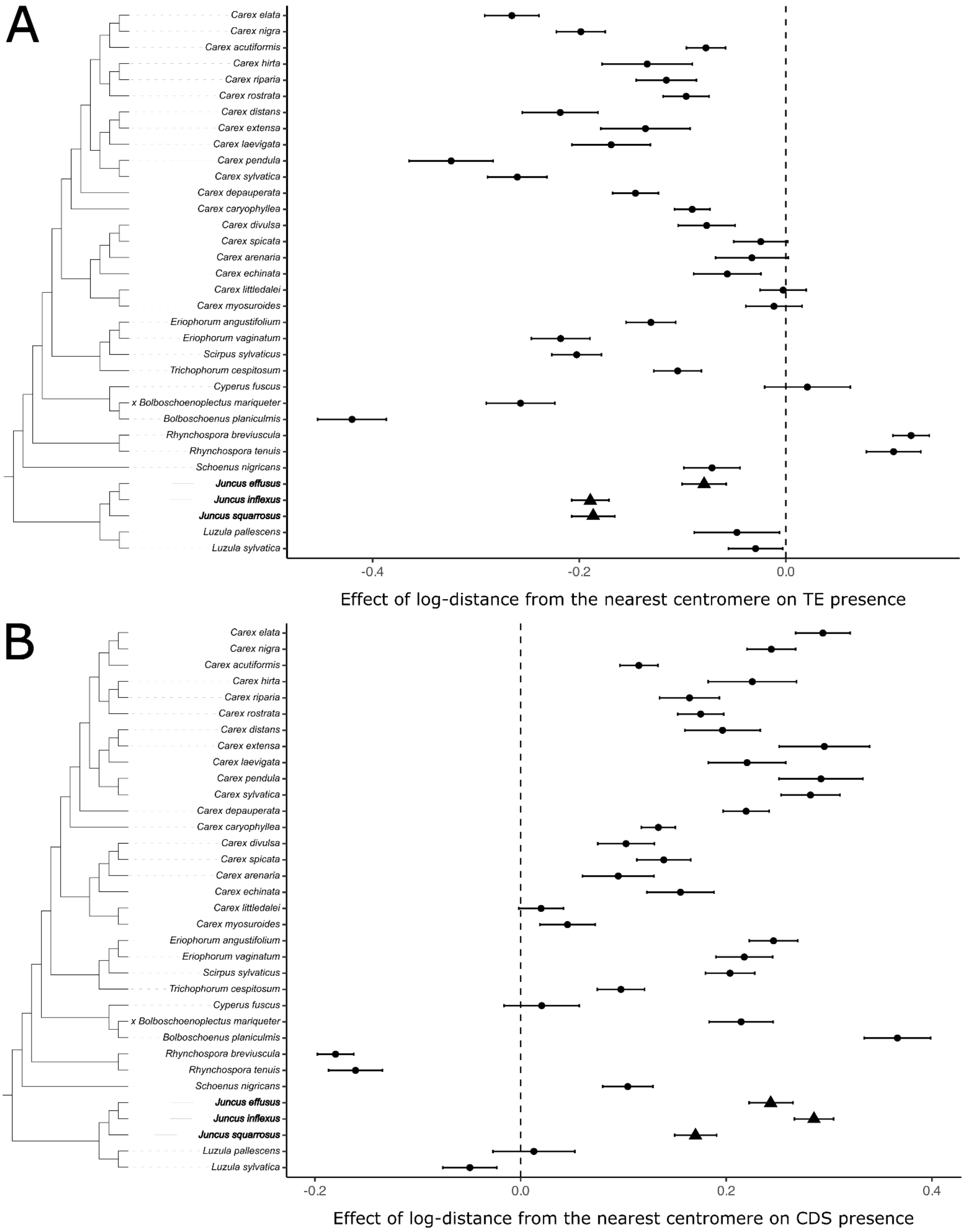


#### **Figure S12**. The effects of polycentromeres on the TE and gene landscape.

**A**. Effect sizes and 95% confidence intervals of a generalised linear model of transposable element overlap with a 5 kb window (identified by RepeatModeler) against distance from the nearest centromere. Monocentric species in bold and indicated by triangles. **B**. As above but for CDS overlap, predicted by Helixer. The phylogenies were made using iTOL (https://itol.embl.de/)[40].
